# Computationally designed sensors for endogenous Ras activity reveal signaling effectors within oncogenic granules

**DOI:** 10.1101/2022.11.22.517009

**Authors:** Jason Z. Zhang, William H. Nguyen, Nathan Greenwood, John C. Rose, Shao-En Ong, Dustin J. Maly, David Baker

## Abstract

Genetically encoded biosensors have accelerated biological discovery, however many important targets such as active Ras (Ras-GTP) are difficult to sense as strategies to match a sensor’s sensitivity to the physiological range of target are lacking. Here, we use computational protein design to generate and optimize intracellular sensors of Ras activity (<u>LOCKR</u>-based <u>S</u>ensor for <u>Ras</u> activity: Ras-LOCKR-S) and proximity labelers of the signaling environment of Ras (<u>LOCKR</u>-based, <u>Ras</u> activity-dependent <u>P</u>roximity <u>L</u>abeler: Ras-LOCKR-PL). We demonstrate that our tools can measure endogenous Ras activity and environment at subcellular resolution. We illustrate the application of these tools by using them to identify Ras effectors, notably Src-Associated in Mitosis 68 kDa protein (SAM68), enriched in oncogenic EML4-Alk granules. Localizing these sensors to these granules revealed that SAM68 enhances Ras activity specifically at the granules, and SAM68 inhibition sensitizes EML4-Alk-driven cancer cells to existing drug therapies, suggesting a possible therapeutic strategy.

## Introduction

Precise modulation of the activity of the signaling enzyme Ras is critical for normal cell function, and Ras is frequently mutated in cancers^1^. Ras GTPases switch between GDP-bound (inactive) and GTP-bound (active) states, and these states are dynamically regulated by guanine exchange factors (GEFs) and Ras GTPase activating proteins (GAPs) that promote the active and inactive states, respectively. Ras-GTP levels can change rapidly in response to external stimuli such as growth factor activation of receptor tyrosine kinases (RTKs) and the dynamics of Ras-GTP levels are based on mechanism of activation and context. Like many intracellular molecules, Ras is dynamically localized to various intracellular organelles through trafficking but its activity and function at particular subcellular regions is not well defined^2–4^. While it was generally thought that Ras requires membranes for activation, membrane-less granules formed by oncoprotein RTK fusions, such as EML4-Alk, have recently been shown to lead to membrane-independent cytosolic Ras activity^5^. However, the mechanisms for activating Ras inside these granules are unclear. Overall, despite decades of study, there are still many open questions about the spatiotemporal activity and regulation of Ras due to the lack of appropriate tools for studying endogenous active Ras (Ras-GTP).

Genetically encoded biosensors are useful for recording the spatiotemporal activities and concentrations of biomolecules in their native context. While optogenetic and chemogenetic systems have been developed that allow direct activation of subcellular populations of Ras, such as Chemically Inducible Activator of Ras (CIAR)^6^, there are no sensors that can measure real-time, sub-cellular activity of endogenous Ras^2,4,7–9^. The lack of sensors for targets like endogenous active Ras is due to the limitations of modular engineering and the grand challenge in matching a sensor’s dynamic range to the physiologically relevant concentration regime of the target^10^ (**Fig. 1a**). Endogenous Ras is believed to only undergo a nanomolar increase in Ras-GTP levels after upstream activation^11^, which makes detection of physiologically-relevant Ras activity difficult. Most methods for generating biosensors involve the modular engineering of native protein domains for detecting and reporting protein targets^12^. The number of suitable native protein domains is limited, and these domains often cannot be engineered without affecting specificity or sensor performance (e.g. ability of sensor to sense target). Thus, only a few variables in sensor development can be altered. Moreover, there are few sensor development methodologies that enable rational tuning of a sensor’s switching and dynamic range^10^ (**Fig. 1a**). To overcome these limitations, we set out to use *de novo* proteins and computational methods to design sensors with switching and dynamic behavior tuned to sense endogenous Ras activity with multiple readouts.

**Figure 1:**
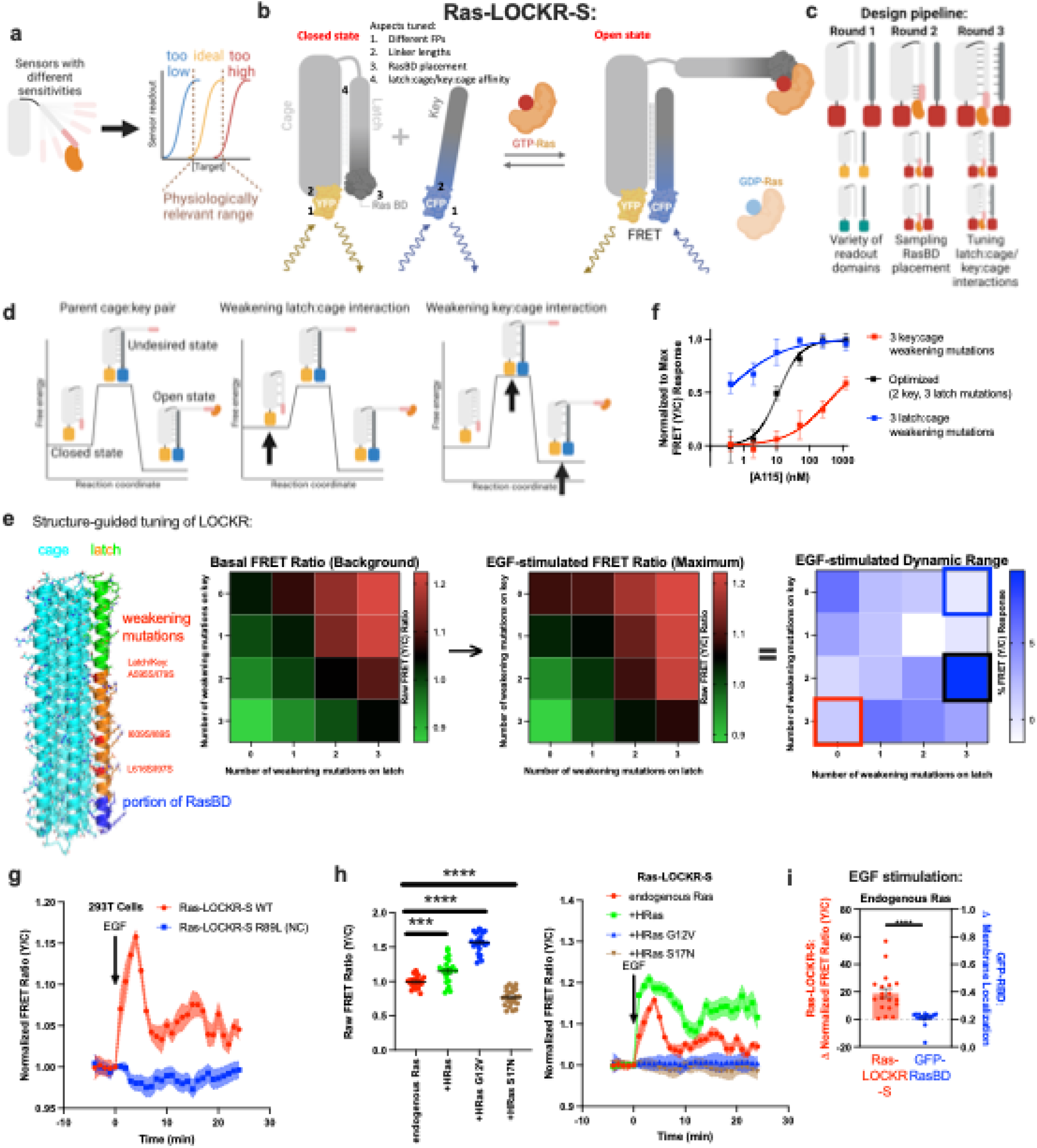
*De novo* designed <u>Ras</u>-reporting <u>LOCKR</u>-based <u>s</u>ensor (Ras-LOCKR-S) measures endogenous Ras activity. **a**, Schematic describing the difficulty in matching sensor’s switching sensitivity to biologically relevant concentration range. **b**, Schematic of optimized Ras-LOCKR-S consisting of fluorescent protein (FP)-tethered Cage and Key where GTP-loaded Ras binds to Ras binding domain (RasBD), thus converting Ras-LOCKR-S from its closed to open state. **c**, Design pipeline for developing Ras-LOCKR-S. **d**, Putative free energy states of Ras-LOCKR-S and how mutations affect its free energy states. **e**, Structure prediction guides the maturation of Ras-LOCKR-S. (left) Predicted structure of Ras-LOCKR-S with mutations highlighted. (right) Heat maps of Ras-LOCKR-S with varying amounts of latch:cage or key:cage weakening mutations. Ras-LOCKR-S mutants were transfected into 293T cells and their yellow/cyan FRET ratio before (background) and after (maximum) 100ng/mL EGF stimulation (n=14 cells per condition). Dynamic range is calculated as maximum FRET ratio after stimulation minus basal FRET ratio. Colored boxes in **e** correspond to the same constructs tested in **f**. **f**, In CIAR-PM-239T cells transfected with Ras-LOCKR-S mutants, varying doses of A115 (n=at least 12 cells per condition). Normalized to maximum FRET responses is calculated by taking the lowest and highest FRET ratios from the entire dataset and normalizing them to 0 and 1, respectively. **g**, Normalized FRET ratio changes in 293T cells transfected with wildtype (WT) or negative control (NC) version of Ras-LOCKR-S and stimulated with 100ng/mL EGF (n=10 cells per condition). **h**, (left) Starting raw FRET ratios and average normalized FRET ratio changes in Ras-LOCKR-S-expressing 293T cells co-expressing nothing else (endogenous Ras), Ras (+HRas), constitutively active Ras (+HRas G12V), or dominant negative Ras (+HRas S17N) (left: n=10 cells per condition, right: n=23 cells per condition). **i**, Comparison of Ras-LOCKR-S to RasBD-GFP^4^ in their response to 100ng/mL EGF in 293T cells (n=19 cells). Solid lines indicate representative average time with error bars representing standard error mean (SEM). Bar graphs represent mean + SEM. ****p < 0.0001, ***p < 0.001, ordinary one-way ANOVA. Scale bars = 10µm.

## Results

### Computational design of LOCKR-based Ras activity sensors

We sought to design Ras sensors with the following properties: 1) sense the activity and environment of Ras, 2) are sensitive enough to detect endogenous levels of active Ras, 3) can be used for live-cell imaging, 4) specifically report events within subcellular regions, 5) and have single cell resolution. We reasoned that the *de novo* designed LOCKR switch system^13,14^ would be well suited to sense active Ras (**Extended Data Fig. 1a**) due to its generalizability, orthogonality, and tunability. LOCKR is a two-component system consisting of a (1) Cage protein that contains a “cage” domain, a “latch” domain that intramolecularly interacts with the cage and embeds a target protein binding domain (TBD), and one portion of a split molecular readout and a (2) Key protein that contains a cage domain-binding “key” domain and the other portion of a split molecular readout. In the absence of a target, the latch of the Cage is closed, and the key domain minimally interacts with the cage domain, resulting in low readout signal. The interaction of a target protein with the TBD located on the latch decreases the latch’s intramolecular interaction with cage, increases the relative intermolecular interaction between key and cage, and allows reconstitution of the two readout portions^13^ (**Fig. 1b**). The LOCKR switch has been demonstrated to allow the measurement of several target molecules *in vitro*^14^ and should be amenable to intracellular applications.

In designing <u>LOCKR</u>-based <u>s</u>ensors for <u>Ras</u> (Ras-LOCKR-S), we split the design process into 3 stages (**Fig. 1c**). In stage 1, we determined the compatibility of the LOCKR system with intracellular readouts (all LOCKR-S candidates and associated experimental results are listed in **Supplementary Table 1**). Previous LOCKR-based sensors have used the reconstitution of split luciferase as the readout of the Key’s intermolecular interaction with the Cage, which is not optimal for subcellular reporting due to its low resolution^12^. Thus, we explored the use of intermolecular, distance-dependent Förster Resonance Energy Transfer (FRET), a commonly used modality for intracellular activity reporters^15^, as a readout for our intracellular LOCKR switches. To do this, we converted the readout of a previously developed LOCKR-based sensor for the receptor binding domain (RBD) of SARS-CoV-2 lucCageRBD^14,16^ to a FRET readout by placing CFP or YFP at the termini of Key or Cage (**Extended Data Fig. 1a-f**). Regardless of the FP placement, RBD addition increased FRET ratios (YFP/CFP) by up to 40% (**Extended Data Fig. 1e**), indicating the feasibility of FRET as a readout for intracellular LOCKR-based sensors.

In stage 2, we engineered the FRET-based LOCKR system to measure endogenous Ras activity by embedding the Ras binding domain (RasBD) from Raf1^17^, which is selective for Ras-GTP, within the latch. To achieve responsiveness to Ras-GTP levels, we computationally designed Ras-LOCKR-S constructs where the embedded RasBD interacts with the cage domain in the closed state (**Fig. 1b, left**). In these designs, binding of Ras-GTP to the embedded RasBD shifts the equilibrium from the closed state with latch bound to cage to the open state (**Fig. 1b, right**) where the Key binds to the Cage, resulting in increased FRET due to the close proximity of the two FPs. To determine the placement of RasBD within the latch, we used the Rosetta-based GraftSwitchMover^13^ algorithm to graft the RasBD onto the C-terminal section of latch at different registers (placements differ by several amino acids, **Supplementary Table 1**). These design efforts identified a subset of seven possible RasBD placements that appeared promising enough to test in 293T cells for their responsiveness to Epidermal growth factor (EGF)-mediated increases in Ras-GTP levels (three designs showed statistically significant increase). The two candidates with the largest FRET ratio increase (1.5%) in response to EGF stimulation also displayed FRET ratio increases upon A115 treatment of 293T cells expressing chemically-induced activator of Ras (CIAR) switch localized to the PM^6^ (CIAR-PM-293T) (**Extended Data Fig 1g**). The highest dynamic range candidate (Ras-LOCKR-S_pos4) from RasBD placement testing was further optimized by ensuring similar expression of Key and Cage using a P2A sequence which improved the EGF-stimulated FRET ratio increase to 5% (**Extended Data Fig. 1h**, all the candidates tested and their results are listed in **Supplementary Table 1**).

In stage 3, we tuned the free energies of the three states of LOCKR-S^14^ (closed, open, and an undesired false positive state where Key binds to Cage without target binding (**Fig. 1d**)) to better match the sensor’s sensitivity to switching within the physiologically relevant range of Ras-GTP (**Fig. 1a**) To do so, we used structure predictions (AlphaFold)^18^ to identify Key and Cage mutations that weaken key:cage and latch:cage interactions (**Extended Data Fig. 1i**) and altered the binding free energies of LOCKR-S‘s three thermodynamic states (**Fig. 1d**). We reasoned that latch:cage-weakening mutations should increase maximum signal and background by allowing greater key:cage interaction, while key:cage-weakening mutations should decrease background and maximum signal by reducing the ability of the Key to compete with the latch for Cage binding. Introducing these mutations into Cages and Keys and testing them in 293T cells, key:cage weakening mutations decreased background and EGF-stimulated FRET ratios while latch:cage weakening mutations increased background and max FRET ratios (**Fig. 1e**), aligning with our predictions.

We evaluated our efforts to tune the sensitivity of Ras-LOCKR-S worked as designed by testing the responses of several candidates with variable key:cage and latch:cage interactions to different levels of Ras-GTP levels, which was achieved by titrating CIAR-PM-293T cells with A115. We found that the Ras-LOCKR-S candidate that possesses a Key with reduced affinity for the Cage (**Fig. 1f**, red line) only responded to high Ras-GTP levels. Meanwhile, the Ras-LOCKR-S candidate with a weakened latch interaction with the cage (**Fig. 1f**, blue line) demonstrated high background FRET at low Ras-GTP levels and a minimal increase in response to higher A115 concentrations (**Fig. 1f**). Ras-LOCKR-S_P2A2_L3K2 (3 mutations in latch and 2 mutations in key) construct showed the highest FRET ratio increase to EGF stimulation (10%) upon EGF stimulation and a robust FRET ratio increase (10%) upon A115 addition in CIAR-PM-293T cells (**Fig. 1e and Extended Data Fig. 1j**). Furthermore, this construct demonstrated the largest range of FRET response across A115 doses (**Fig. 1f**, black line, “optimized”), showcasing that it is well suited for sensing the physiological range of endogenous Ras-GTP levels; this is the Ras-LOCKR-S construct used throughout the rest of the paper. These results demonstrate that Ras-LOCKR-S can measure a range of endogenous Ras activities, and that the LOCKR system is an ideal platform for rationally designing intracellular sensors that are specifically tuned for intracellular applications of interest.

We characterized the temporal response of a Ras-LOCKR-S construct that is not targeted to any subcellular region in response to EGF stimulation. Consistent with the well-characterized dynamics of Ras-GTP levels that result from EGF stimulation, we observed a rapid and transient increase in FRET ratios, demonstrating Ras-LOCKR-S’s rapid sensing and reversibility in response to variable Ras-GTP levels. We found that Ras-LOCKR-S provided sustained increases in FRET ratios after treating CIAR-PM-239T cells with A115 (**Fig. 1g and Extended Data Fig. 1k**), reflecting the dynamics previously observed with Ras-GTP pulldowns and phospho-Erk analysis^6^. Under the same conditions, we found that there were no changes in FRET ratios for a negative control Ras-LOCKR-S construct (NC) that contains a mutation (RasBD^R89L^) that abrogates Ras-GTP binding^17^, indicating that the FRET response from Ras-LOCKR-S is due to Ras-GTP binding. Further confirming Ras-LOCKR-S’s specificity for sensing Ras-GTP, we observed that co-expression of either a constitutively active mutant of Ras (Hras^G12V^) or a dominant negative mutant of Ras (Hras^S17N^) increased and decreased raw FRET ratios, respectively (**Fig. 1h**). Co-expression of Hras^S17N^ also eliminated FRET ratio changes after EGF stimulation or A115 activation of CIAR-PM-239T cells (**Fig. 1h and Extended Data Fig. 1l**). Furthermore, co-expression of a GAP for Rap GTPase (structurally similar to Ras) minimally influenced Ras-LOCKR’s response to EGF stimulation (**Extended Data Fig. 1m**) but did abrogate a Rap1 biosensor’s (Rap1 FLARE)^19^ response to EGF (**Extended Data Fig. 1n**) In comparing to the commonly used GFP-RasBD^4^ construct commonly used to measure Ras activity, only Ras-LOCKR-S can report endogenous Ras activation from EGF stimulation (**Fig. 1i**). Overall, Ras-LOCKR-S is a quantitative sensor capable of real-time readout of endogenous Ras activity.

### Ras-LOCKR-S detects endogenous Ras activity in different subcellular compartments and in multiple cell types

Next, we developed subcellularly-localized Ras-LOCKR-S to map out the subcellular landscape of endogenous Ras signaling (**Fig. 2a**). We pursued a strategy that restricted the Key to a subcellular region of interest to increase the likelihood that the signals detected (and thus Ras activities) are emanating specifically from that particular area^20,21^. We localized the Key of Ras-LOCKR-S to the PM or Golgi using localization tags (PM: N-terminus of Lyn^22^, Golgi: N-terminus of eNOS^23^) (**Fig. 2b**). While we observed that the cage was diffusely localized, the Key was restricted to the intended subcellular compartment (**Fig. 2b**). Furthermore, FRET signals primarily emanated from the targeted subcellular regions in response to EGF stimulation (**Fig. 2c-d**). EGF or A115 addition led to FRET ratio increases primarily at the PM for PM localized Ras-LOCKR-S (PM-Ras-LOCKR-S) (**Fig. 2c**). We next explored if the Golgi-localized Ras-LOCKR-S (Golgi-Ras-LOCKR-S) could detect endogenous Ras-GTP at the Golgi (**Extended Data Fig. 2a**) by directly activating Ras at this subcellular location with a Golgi-localized CIAR (CIAR-Golgi) construct. We found that addition of A115 in CIAR-Golgi-expressing 293T cells led to an increase in FRET ratios for Golgi-Ras-LOCKR-S but not for PM-Ras-LOCKR-S (**Extended Data Fig. 2b**), confirming that these localized Ras sensors are reporting compartment specific activities. We also found that both EGF and A115 addition in CIAR-PM-293T cells led to FRET ratio increases for Golgi-Ras-LOCKR-S (**Fig. 2d**), demonstrating that endogenous Ras activity indeed occurs at endomembranes presumably due to Ras trafficking from the PM to endomembranes^24^. Untargeted and targeted Ras-LOCKR-S expressed at similar levels (**Extended Data Fig. 2c**) and sensor expression levels did not obviously correlate with their EGF-stimulated FRET ratio increases (**Extended Data Fig. 2d**), thus the expression levels of different localized Ras-LOCKR-S do not account for these functional differences. Overall, localized Ras-LOCKR-S can detect subcellular Ras activation levels and provide insight into subcellular Ras signaling networks.

**Figure 2:**
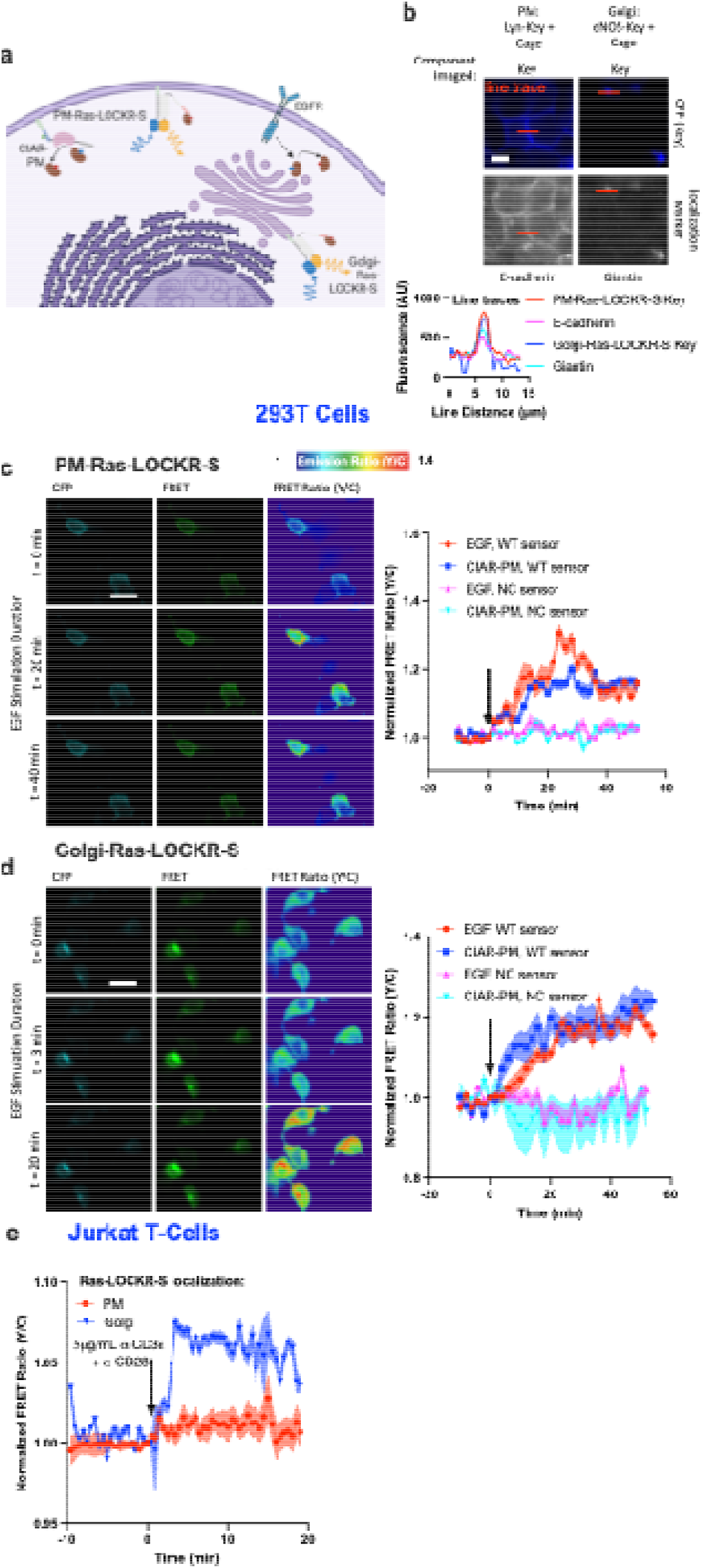
Ras-LOCKR-S can report subcellular endogenous Ras activities in various cell types. **a**, Schematic of subcellularly targeted Ras-LOCKR-S, EGF receptor (EGF), and CIAR-PM. **b**, (top) Representative images of 293T cells transfected with localized Ras-LOCKR-S (Key with localization sequence, Cage untargeted) and stained for established localization markers. (bottom) Line trace comparison of CFP signal from Ras-LOCKR-S Key and localization marker. **c-d**, CIAR-PM-239T cells expressing Ras-LOCKR-S WT or NC localized to PM (**c**) or Golgi (**d**) were stimulated either with 100ng/mL EGF or 250nM A115. (left) Representative epifluorescence images (CFP channel from Key, FRET channel, and pseudocolored raw FRET ratio). (right) Normalized FRET ratio changes (right, n=at least 15 cells per condition). **e**, Average normalized FRET ratio responses of Jurkat T-cells expressing Ras-LOCKR-S localized to Golgi or PM stimulated with 5µg/mL of a-CD3E + a-CD28 (n=at least 10 cells per condition). Solid lines in **c-d** indicate a representative average time course with error bars representing standard error mean (SEM). Solid lines in **e** indicate average time courses of FRET ratio changes from all cells combined from 3 experiments with error bars representing standard error mean (SEM).

Next, we explored the utility of Ras-LOCKR-S to report subcellular endogenous Ras activities in other cell types. Compared to 293T cells, Ras in Jurkat T-cells is highly enriched at the Golgi (**Extended Data Fig. 2e**)^25,26^. We found that T-cell receptor (TCR)-activated Jurkats displayed FRET ratio increases for Golgi-Ras-LOCKR-S but not PM-Ras-LOCKR-S (**Fig. 2e**), which demonstrates that TCR activation leads to endogenous Ras activity at the Golgi. This observation is consistent with a study that shows TCR stimulation activates Ras at the Golgi in Jurkats overexpressing NRas, which is primarily at the Golgi, and aligns with a report showing that co-stimulation of the TCR and lymphocyte function-associated antigen-1 is required for Ras signaling at the PM^25,26^. These Jurkat results demonstrate that Ras-LOCKR-S can report endogenous Ras activity in a different cell type and that targeted Ras-LOCKR-S reports compartment-specific Ras activities.

### Computational design of LOCKR-based Ras activity-dependent proximity labelers

To enable identification of upstream Ras activators and downstream effectors in different cellular compartments, we developed a LOCKR-based system that carries out proximity labeling in response to Ras-GTP (<u>LOCKR</u>-based, <u>Ras</u>-activity dependent <u>p</u>roximity labeler: Ras-LOCKR-PL) (**Fig. 3a and Supplementary Table 2**, which lists all LOCKR-PL candidates and their results). We reasoned that such a Ras-GTP-responsive proximity labeler could be generated by replacing the FPs in Ras-LOCKR-S with split biotin ligases (**Fig. 1c**), allowing assembly of a functional proximity labeler upon binding of Ras-GTP to RasBD (**Fig. 3a**).

**Figure 3:**
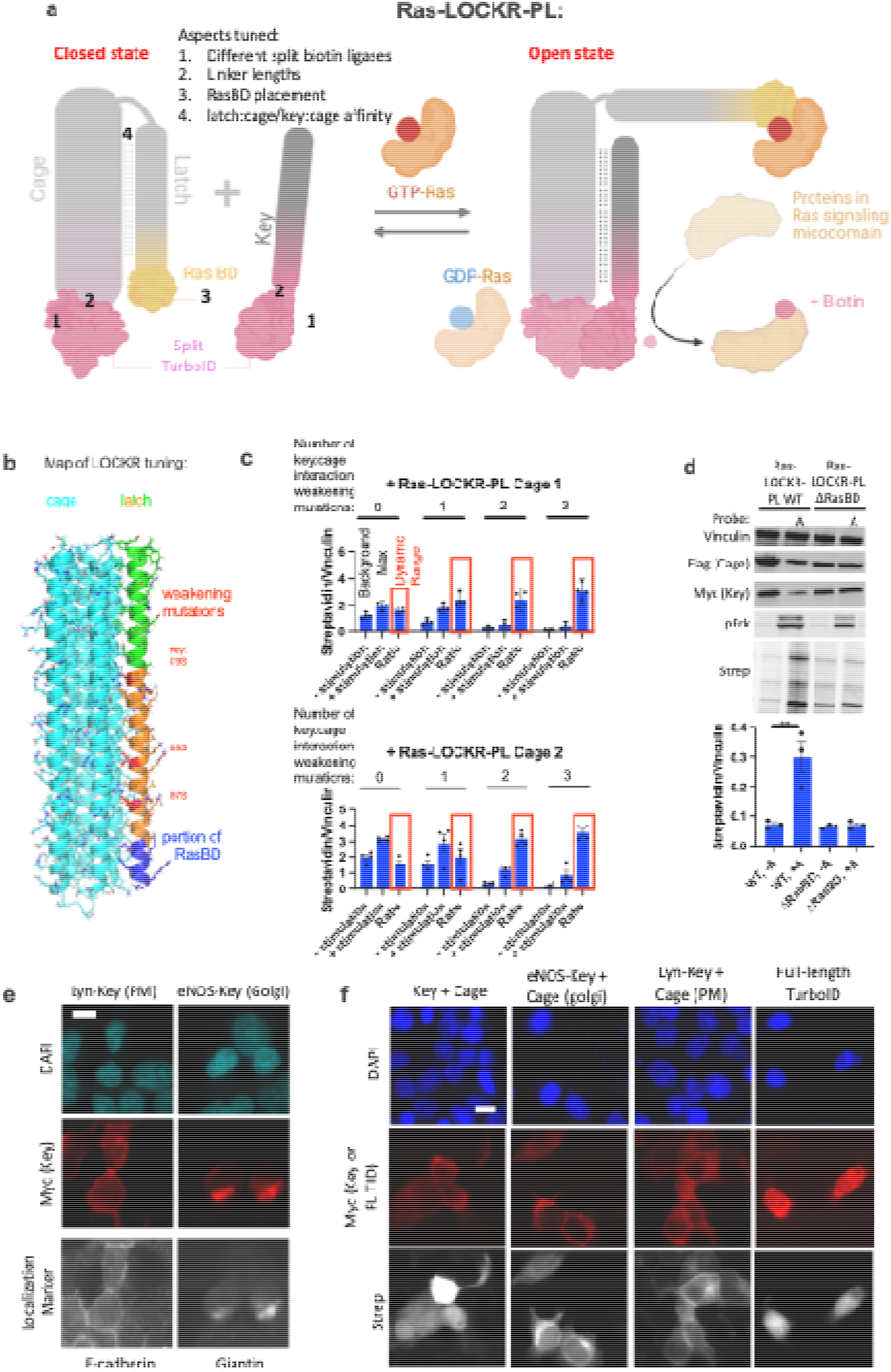
*De novo* designed <u>Ras</u>-dependent <u>LOCKR</u>-based <u>p</u>roximity labeler (Ras-LOCKR-PL) profiles components in specified Ras signaling microdomains. **a,** Schematic of optimized Ras-LOCKR-PL consisting of split TurboID tethered to Cage and Key where GTP-Ras binds to RasBD, allowing the reconstitution of functional TurboID and thus biotinylation of neighboring proteins. **b**, Predicted structure of Ras-LOCKR-PL with mutations highlighted. **c**, Bar graph of CIAR-PM-239T cells were transfected with Ras-LOCKR-PL candidates and 500 M biotin was added for 16 hours without (-stimulation) or with (+stimulation) 250nM A115 (n=4 experiments per condition). Ratio is +stimulation divided by -stimulation. **d**, Representative immunoblots of CIAR-PM-239T cells transfected with Ras-LOCKR-PL WT or a mutant without RasBD (ΔRasBD) with 500 M biotin with or without 250nM A115 (labeled “A”) for 16 hours (n=3 experimental repeats). **e-f**, Subcellularly localized Ras-LOCKR-PL or full-length TurboID expressed and underwent immunostaining with antibodies for established localization markers or streptavidin. Bar graphs represent mean + SEM. **p < 0.01 unpaired two-tailed Student’s t-test. Scale bars = 10µm.

Similar to our design strategy for Ras-LOCKR-S (**Fig. 1c**), for stage 1 of constructing Ras-LOCKR-PL, we tested whether split versions of the biotin ligases ContactID^27^ and TurboID^28^ can be incorporated into the LOCKR system by swapping the split luciferase in lucCageRBD^14,16^ with split biotin ligase (RBD-LOCKR-PL). In 293T cells expressing RBD, the split ContactID-containing RBD-LOCKR-PL (**Extended Data Fig. 3a-b**) showed no Increase in biotinylation of cellular proteins, while the split TurboID-containing RBD-LOCKR-PL (**Extended Data Fig. 3c-d**) showed RBD-dependent biotinylation. Thus, we used split TurboID for constructing an optimized Ras-LOCKR-PL.

For stage 2 of our Ras-LOCKR-PL design, we again used the SwitchGraftMover algorithm to identify possible RasBD placements on latch. Several of the most promising rationally selected candidates were tested for Ras activity-induced labeling in CIAR-PM-293T cells treated with A115 for 16 hours (**Extended Data Fig. 3e-g**). We found that an offset of five amino acids relative to the RasBD placement for Ras-LOCKR-S led to a Ras-LOCKR-PL construct with the highest dynamic range (100% increase in biotinylation in A115-treated conditions) (**Extended Data Fig. 3f**).

For stage 3 of Ras-LOCKR-PL design, we again used protein structure prediction models to find mutations or truncations of the latch and key to tune their interactions with cage to maximize the dynamic range in response to A115 treatment of CIAR-PM-293T cells (**Fig 3b**). Mutating the latch to decrease the latch:cage interaction strength increased background but not A115-stimulated signal, resulting in Ras-LOCKR-PL candidates with a decreased dynamic range (**Extended Data Fig. 3h**). In contrast, mutations that weaken the key:cage interaction consistently decreased background biotinylation, leading to Ras-LOCKR-PL candidates with an overall increased dynamic range (200-300% increase in biotinylation in A115-treated conditions) (**Fig. 3c and Extended Data Fig. 3i**). As expected, weakening of the key:cage interaction led to a proportional decrease in background biotinylation (**Fig. 3c and Extended Data Fig. 3i**).

The highest dynamic range Ras-LOCKR-PL candidates (boxed in green in **Extended Data Fig. 3i**) were further tested for speed and accuracy at shorter biotinylation times (1 and 3 hours) with A115 treatment or EGF stimulation in CIAR-PM-293T cells (**Extended Data Fig. 3j**). Of the designs that showed increased biotinylation within 3 hours of Ras activation, Ras-LOCKR-PL_Km3C2 (no weakening mutations in latch and 3 weakening mutations in key) showed consistent biotin labeling of Ras (**Extended Data Fig. 4a**); we refer to this design as Ras-LOCKR-PL for the remainder of the paper. This optimized Key and Cage for Ras-LOCKR-PL differs from Ras-LOCKR-S in requiring a tighter latch:cage interaction to prevent background signal, which is likely due to the innate affinity between the two halves of split TurboID (unlike YFP and CFP) and the hours-long labeling time required. To further validate that labeling by Ras-LOCKR-PL is dependent on active Ras, we co-expressed Ras-LOCKR-PL with constitutively active Ras (Hras^G12V^) or the catalytic domain of Sos (Sos_cat_) in CIAR-PM-239T cells treated with or without A115 for 16 hours. Co-expression of these factors led to increased levels of biotin labeling and phospho-Erk levels even without A115 addition. In contrast, co-expression of dominant negative Ras (Hras^S17N^) or the RasGAP Gap1m blunted biotinylation and pErk even with A115 (**Extended Data Fig. 4b**). Deletion of the RasBD in Ras-LOCKR-PL eliminated A115-induced biotinylation in CIAR-PM-239T cells (**Fig. 3d**), demonstrating that Ras-LOCKR-PL relies on Ras-GTP binding to induce biotinylation. Further confirming that labeling by Ras-LOCKR-PL is Ras-GTP specific, co-expression of a RapGEF (CalDAG-GEFI) to increase Rap-GTP levels^9^ did not affect A115-induced biotinylation (**Extended Data Fig. 4b**). Altogether, these validation experiments demonstrate that Ras-LOCKR-PL is a Ras activity dependent proximity labeler.

We next sought to subcellularly target the Key of Ras-LOCKR-PL using the same localization sequences as Ras-LOCKR-S and observed that each construct colocalized with the expected subcellular marker by immunofluorescence (**Fig. 3e**). Functionally, biotinylation by localized Ras-LOCKR-PL in CIAR-PM-239T cells treated with A115 for 3 hours was spatially restricted to expected subcellular regions (**Fig. 3f and Extended Data Fig. 4c**). Furthermore, we found that the subcellularly localized Ras-LOCKR-PL system does not affect global (**Extended Data Fig. 3i and 4c**) or compartment-specific downstream Erk activity, as measured by subcellularly localized EKAR4^29^ (**Extended Data Fig. 4d-e**), suggesting that Ras-LOCKR-PL does not significantly buffer signaling downstream of Ras-GTP. Overall, Ras-LOCKR-PL is a Ras signaling-dependent proximity labeler that accurately profiles active subcellular signalosomes (Ras-GTP hotspots). Moreover, successful construction of two sensors that have different readouts (fluorescence, proximity labeling) for the same target (endogenous Ras-GTP) demonstrates the generality of this sensor scaffold and design strategy.

### Ras sensors identify components inside Ras-active EML4-Alk granules that drive aberrant cytosolic Ras signaling

In addition to membranes, Ras signaling can occur in membrane-less oncogenic granules, like granules formed by EML4-Alk fusions, which are found in 2-9% of non-small-cell lung cancers^30^. EML4-Alk variant 1 (v1) and variant 3 (v3), which represent the majority of observed oncogenic fusions, form cytosolic granules that can recruit upstream activators of Ras via the Alk kinase domain^5,31^. We explored if our Ras-LOCKR-S can be used to measure Ras activity in EML4-Alk granules. To allow the sensing of Ras activity inside granules, the Key of Ras-LOCKR-S was fused to EML4-Alk, which was co-expressed in Beas2B lung cells with an untargeted Cage. We found that fusing the Key to EML4-Alk did not affect granule formation (**Fig. 4a**). Consistent with previous reports suggesting that increased Ras activity exists within EML4-Alk-containing granules^5^, we observed increased raw FRET ratios in punctate regions compared to the diffuse regions of Beas2B cells expressing the EML4-Alk targeted Ras-LOCKR-S system (**Fig. 4a and Extended Data Fig. 5a**). In contrast, FRET ratios were low in punctate and diffuse regions in cells expressing NC Ras-LOCKR-S (**Extended Data Fig. 5a**). Furthermore, when the Key was fused to a EML4-Alk variant that does not form granules (Δ Trimerization Domain (TD))^5,31^, similarly low levels of FRET were observed (**Extended Data Fig. 5a**). Thus, Ras-LOCKR-S can measure endogenous Ras activity in a variety of subcellular compartments, including membrane-less granules.

**Figure 4:**
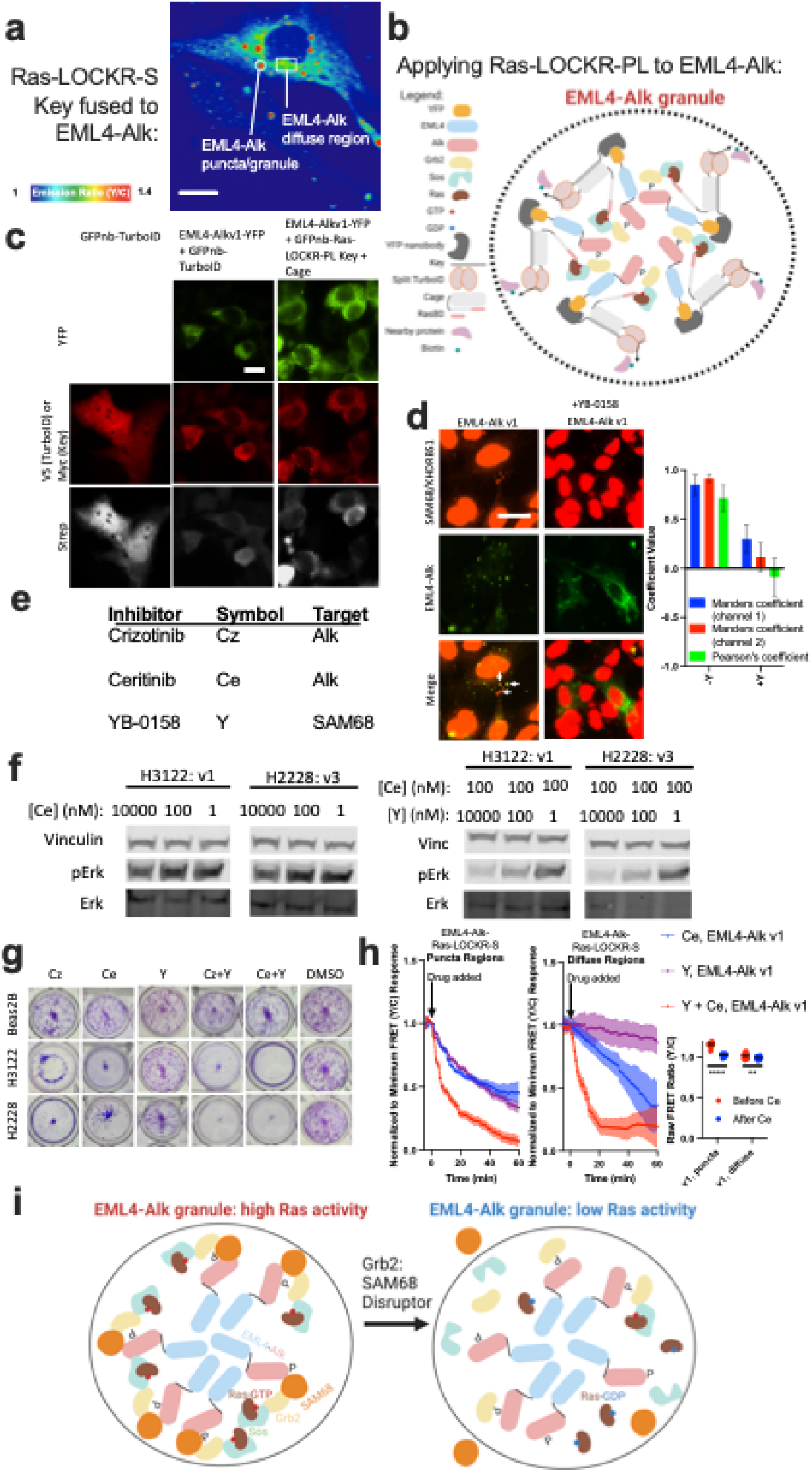
Identifying upstream drivers of oncogenic Ras activity inside EML4-Alk granules. **a**, Representative pseudocolored FRET ratio image of Beas2B cell transfected with Ras-LOCKR-S localized to EML4-Alk (Key fused to EML4-Alk, Cage untargeted). **b**, Schematic of strategy for localizing Ras-LOCKR-PL to EML4-Alk granules. **c**, Representative epifluorescence images of Beas2B cells transfected with YFP-fused EML4-Alk variant 1 (v1) and GFP nanobody (GFPnb)-fused V5-tagged TurboID or GFPnb-fused Myc-tagged Ras-LOCKR-PL. 500µM biotin was added to cells for 3 hours and then stained for streptavidin labeling. **d**, Left: Representative epifluorescence images of Beas2B cells expressing YFP-tagged EML4-Alk v1, treated with 1µM YB-0158 for 1 hr and immunostained for SAM68. Arrows indicate co-localization of EML4-Alk with SAM68 puncta. Right: colocalization analysis. **e**, Table of inhibitors used in this figure. **f**, Representative pErk immunoblots of H3122 (EML4-Alk v1 positive) or H2228 (EML4-Alk v3 positive) cancer patient cells treated with indicated concentrations of inhibitors. **g**, Representative images of crystal violet staining of Beas2B, H3122, and H2228 incubated for 1 week with inhibitors to Alk (Cz: 1µM Crizotinib, Ce: 1µM Ceritinib), SAM68 (Y: 1mM YB-0158), or DMSO. **h**, (left) Normalized to minimum FRET ratio time-courses of Beas2B cells transfected with Ras-LOCKR-S localized to EML4-Alk v1 and incubated with 10µM of inhibitors. Puncta and diffuse regions were analyzed separately. Normalized to minimum FRET ratios are calculated by normalizing the data set to the condition with the largest decrease in FRET ratios (Ce + Y in both cases) where 0 represents the lowest FRET ratio out of the entire data set. (right) Raw FRET ratios of puncta and diffuse EML4-Alk regions after Alk inhibition for 1 hour. **i**, Schematic of how SAM68 inhibition by disrupting SAM68:Grb2 interactions decreases local Ras activity in EML4-Alk granules. Bar graphs represent mean + SEM. ****p < 0.0001, **p < 0.01 one-way ANOVA. Scale bars = 10µm.

Several known Ras effectors have been shown to be located within EML4-Alk granules^5^, but an unbiased investigation of the membrane-less Ras signalosome has not been performed due to lack of appropriate tools. To provide insight into the signaling environment of active Ras within granules, we generated granule-localized versions of Ras-LOCKR-PL (**Fig. 4b**). Granule formation was lost when the Key of Ras-LOCKR-PL was fused to EML4-Alk (**Extended Data Fig. 5b**) but retained when the Key was fused to a GFP nanobody (GFPnb) that targets EML4-Alk-YFP (**Fig. 4b-c and Extended Data Fig. 5c-d**). Consistent with increased Ras signaling inside EML4-Alk granules, Ras-LOCKR-PL recruitment to Ras-active granules resulted in substantial biotinylation inside the granule relative to outside the granule (**Fig. 4c and Extended Data Fig. 5d**). Distinguishable biotinylation within EML4-Alk-containing granules was not observed in cells expressing full length TurboID fused to a GFPnb (**Figure 4c and Extended Data Fig. 5d**), highlighting the unique ability of Ras-LOCKR-PL to selectively label proteins within Ras signaling microdomains.

To broadly profile the Ras signalosome inside EML4-Alk-containing granules, we performed first a set of mass spectrometry (MS) experiments comparing labeled proteins in Beas2B cells co-expressing EML4-Alk v1 and GFPnb-Ras-LOCKR-PL versus cells expressing GFPnb-Ras-LOCKR-PL alone, thus using the same proximity labeling system (GFPnb-Ras-LOCKR-PL) and comparing between with or without EML4-Alk expression (**Extended Data Fig. 5e-f, and Supplementary Table 3**). We identified a number of proteins that were selectively enriched in EML4-Alk v1-expressing cells after 3 or 16 hours of biotin labeling. For further analysis, we focused on proteins that were selectively labeled (more than 2-fold change) in EML4-Alk v1-expressing cells, passed our statistical cutoff, and were related to signaling based on gene ontology analysis (see **Extended Data Fig. 5f** figure legend for details). Proteins that demonstrated increased labeling in Beas2B cells expressing EML4-Alk v1 granules include previously characterized EML4 interactors, like tubulin^31^ (TUBA1B, TUBB, TUBB3, TUBB4B, TUBB8) (blue arrows in **Extended Data Fig. 5f**), and known effectors of the RTK/Ras/MAPK pathway^5,31^, like Grb2, MAP2K1/MEK, and PTPN11/Shp2 (blue arrows in **Extended Data Fig. 5f**). In contrast, Akt, which was previously shown to be excluded from EML4-Alk granules^5^, was found to exhibit reduced labeling in Beas2B cells expressing EML4-Alk fusions than in those that do not (red arrow in **Extended Data Fig. 5f**) and in subsequent MS experiments as well (**Extended Data Fig. 5g-h**). These results demonstrate the ability of Ras-LOCKR-PL to identify components of active Ras signalosomes. To validate that these proteins are indeed enriched in EML4-Alk v1 granules, we performed a second set of MS experiments in EML4-Alk v1-expressing Beas2B cells comparing between targeted Ras-LOCKR-PL (GFPnb-Ras-LOCKR-PL) vs untargeted Ras-LOCKR-PL to identify components selectively enriched (>2-fold change) in EML4-Alk granules (EML4-Alk v1 and GFPnb-Ras-LOCKR-PL versus EML4-Alk v1 and Ras-LOCKR-PL) (**Extended Data Fig. 5g-h and Supplementary Table 3**). The components identified in the first set of MS experiments were also seen to be selectively enriched inside EML4-Alk v1 granules in the subsequent set of MS experiments.

We next validated proteins identified in the first set of MS inside EML4-Alk granules that were not characterized before to be involved in these granules. 14-3-3 proteins (YWHAG, YWHAZ, YWHAE, YWHAB) are well established to interact with phosphorylated components of the MAPK pathway^32^, the cascade downstream of Ras signaling, and were also selectively labeled in EML4-Alk v1-expressing Beas2B cells (black arrows in **Extended Data Fig. 5f**). The second set of MS experiments (comparing EML4-Alk v1 with targeted or untargeted Ras-LOCKR-PL) consistently showed increased labeling of 14-3-3 proteins in EML4-Alk v1 granules (**Extended Data Fig. 5g-h**). To validate these findings, Beas2B cells were transfected with YFP-tagged EML4-Alk v1 and immunostaining of these cells confirmed the enrichment of YWHAG in EML4-Alk v1 granules (**Extended Data Fig. 6a**). In Beas2B cells ectopically expressing V5-tagged EML4-Alk v1, EML4-Alk was immunoprecipitated via its V5 tag and YWHAG (and most likely other 14-3-3 proteins) was probed (**Extended Data Fig. 6b**). YWHAG co-immunoprecipitated with EML4-Alk v1 and YWHAG lost co-immunoprecipitation when the ΔTD mutant of EML4-Alk was expressed which loses granule formation^5,31^ (**Extended Data Fig. 6b**), suggesting that YWHAG (and likely other 14-3-3 proteins) is enriched in EML4-Alk granules specifically. A regulator for Rho GTPase (RhoGDI)^33,34^ was also selectively labeled in the first set (black arrows in **Extended Data Fig. 5e-f**) and second set of MS experiments (**Extended Data Fig. 5g-h**). Furthermore, proteins known to interact with RhoGDI, such as Cdc42, based on STRING databases were also enriched in EML4-Alk v1-expressing Beas2B cells (gray arrows in **Extended Data Fig. 5f and h**). To again validate these findings, immunostaining of YFP-EML4-Alk v1-expressing Beas2B cells confirmed the enrichment of RhoGDI in EML4-Alk v1 granules (**Extended Data Fig. 6a**). RhoGDI co-immunoprecipitated with V5-tagged EML4-Alk v1-expressing in Beas2B cells while co-immunoprecipitation was lost in V5-tagged EML4-Alk v1 ΔTD mutant (**Extended Data Fig. 6b**), suggesting that RhoGDI is selectively enriched in EML4-Alk granules. Thus, these proximity labeling tools can identify new components inside these oncogenic granules.

We further explored the function of Src-Associated in Mitosis 68 kDa protein (SAM68/KHDRBS1), which demonstrated higher labeling in EML4-Alk v1 and v3 granules (**Extended Data Fig. 5e-h**), because it has previously been characterized to interact with the Ras effector Grb2^35^ (also enriched in EML4-Alk-expressing Beas2B cells (**Extended Data Fig. 5e-h**)) and the correlation of its cytosolic localization with tumor grade of renal cell carcinoma^36^. Indeed, the cytosolic pool of SAM68 was enriched in EML4-Alk v1 granules based on immunostaining of Beas2B cells transfected with YFP-tagged EML4-Alk v1 (**Fig. 4d**, note that SAM68 is an RNA-binding protein and is mostly localized in the nucleus^36^). SAM68 also co-immunoprecipitated with EML4-Alk v1 and lost its co-immunoprecipitation with the ΔTD mutant^5,31^ of EML4-Alk (**Extended Data Fig. 7a**), suggesting that SAM68 is indeed sequestered preferentially to EML4-Alk granules. We next explored if inhibition of SAM68 by the small molecule YB-0158^37^ (Y) (**Fig. 4e**), which disrupts SAM68 binding to Grb2 in Beas2B cells (**Extended Data Fig. 7b-c**), blocks EML4-Alk fusion-driven oncogenic signaling and proliferation in patient-derived EML4-Alk cancer cells: H3122 (EML4-Alk v1) and H2228 (EML4-Alk v3). We observed that SAM68 inhibition (Y) minimally diminished pErk levels at all concentrations tested (**Fig. 4f and Extended Data Fig. 7d**). Aligning with previous studies^31^, various concentrations of Alk inhibitor (Ce) for 1 hour led only to a slight decrease in pErk levels (**Fig. 4f**). However, YB-0158 dose dependently decreased pErk levels in the presence of a fixed concentration (100nM) of Alk inhibitor (**Fig. 4f**) and a fixed concentration of YB-0158 (100nM) facilitated inhibition of pErk by Ce (**Extended Data Fig. 7e**), suggesting that SAM68 contributes to Ras-driven MAPK signaling. Co-inhibition of SAM68 and Alk dose dependently (changing doses of both inhibitors simultaneously) decreased pErk levels and led to more potent inhibition of Erk phosphorylation compared to Alk or SAM68 inhibitor alone (**Extended Data Fig. 7d**). These inhibitors dose dependently (changing doses of both inhibitors simultaneously) inhibited cell growth of EML4-Alk addicted cancer cells but not WT lung cells (compared to Alk or SAM68 inhibitor alone) (**Extended Data Fig. 7f**). Co-inhibition also decreased colony formation (**Fig. 4g**) more than single drug treatment and this effect was specific to EML4-Alk-containing cells, showcasing the potential of this drug combination as a potential therapeutic for EML4-Alk driven cancers.

We next used Ras-LOCKR-S to provide more insight into how co-inhibition of Alk and SAM68 leads to reduced Erk signaling and inhibited cell growth. One possible mechanism is that SAM68 potentiates Ras signaling specifically within EML4-Alk granules (see Discussion), thus we probed the effects on subcellular Ras activity when disrupting SAM68:Grb2 interaction by YB-0158 using EML4-Alk-targeted Ras-LOCKR-S (**Fig. 4a**). SAM68 inhibitor YB-0158 (Y) decreased SAM68 enrichment in EML4-Alk granules but did not dissolve the granules themselves (**Fig. 4d**), demonstrating that SAM68 is not involved in granule formation. Performing time-course imaging in Beas2B cells transfected with EML4-Alk-targeted Ras-LOCKR-S, we observed that 1 hour acute incubation with Alk inhibitor (Ce) alone decreased Ras activity inside EML4-Alk v1 and v3 granules (using EML4-Alk-tethered to Ras-LOCKR-S Key) (∼-50%, FRET ratio is normalized to the drug stimulation with maximum decrease in the puncta, see figure legend) but did not dissolve granules (**Fig. 4h and Extended Data Fig. 7g**). As EML4-Alk has both a punctate and diffuse pool, we wondered whether Alk inhibition also decreased Ras activity outside the granule (diffuse regions) (**Fig. 4a**). Although the diffuse regions had lower basal levels of Ras activity (**Fig. 4h and Extended Data Fig. 7g, right**), Alk inhibition indeed led to a decrease in Ras activity in the diffuse regions (∼-60%, FRET ratio is normalized to the drug stimulation with maximum decrease in the diffuse region, see figure legend) (**Fig. 4h and Extended Data Fig. 7g**), suggesting that EML4-Alk in the diffuse regions still signals to Ras. In contrast, SAM68 inhibition decreased Ras activity primarily within the granule (∼-65%) but not in the diffuse regions (∼-5%) (**Fig. 4h and Extended Data Fig. 7g-h**), suggesting that SAM68 distinctly enhances Ras activity exclusively within the granules. Co-inhibition of Alk and SAM68 led to the most dramatic decrease in Ras activities in both the punctate (∼-90%) and diffuse (∼-80%) regions (**Fig. 4h and Extended Data Fig. 7g**) but did not disrupt EML4-Alk granules (**Extended Data Fig. 7i**), demonstrating the utility of this co-inhibition regimen in ensuring blockage of oncogenic Ras activity. Overall, deployment of our Ras-LOCKR tools identify SAM68 as an upstream mediator of Ras activity during EML4-Alk signaling and its mode of action by revealing the subcellular function of SAM68 inhibition (**Fig. 4i**). Translationally, these results suggest the benefit of using SAM68 and Alk inhibitor combination to block oncogenic signaling within EML4-Alk-addicted cancers.

## Discussion

Genetically encoded biosensors are powerful tools for measuring the spatiotemporal dynamics of targets inside their natural environments^15^. Rational design of these sensors is complicated by limitations of current engineering platforms and methods for translating a binding event or activity event into a readout. Moreover, rationally designing sensors where changes in the readout match the biologically relevant concentration range of target is particularly challenging. Here, we show that the *de novo* designed LOCKR protein switch and computational design methodologies enable matching sensor switching with the physiologically relevant range of a target (here, endogenous Ras-GTP) that has eluded previous biosensor design efforts (**Fig. 1a**). We design sensors for measuring real-time, subcellular endogenous Ras activity (Ras-LOCKR-S) at the single cell level and for profiling the interactome of Ras-GTP using proximity labeling (Ras-LOCKR-PL). Ras-LOCKR-PL is particularly useful as it profiles Ras “signalosomes”^38–40^, which are signaling hotspots or microdomains where unique biochemical activities occur and thus mediate specific functions. Both sensors can be subcellularly localized to measure compartment-specific endogenous Ras activities and environment.

Using these new sensors, we found that endogenous Ras is active not only at the PM but also at endomembranes (Golgi) and in membrane-less, cytosolic granules and gained insight into mechanisms of oncogenic signaling. Targeting our Ras-LOCKR tools to EML4-Alk-containing granules enabled the discovery of several unanticipated factors inside these granules. We identified SAM68 as an upstream effector of aberrant cytosolic Ras activity inside EML4-Alk granules, thus providing a more complete picture for how Ras can be activated and is capable of driving oncogenic signaling in the absence of membranes. As a small molecule that inhibits SAM68:Grb2 interaction (**Extended Data Fig. 6e**) disrupts SAM68 sequestration into EML4-Alk granules (**Fig. 4d**), we speculate that SAM68 sequestration into EML4-Alk granules may involve SAM68’s several proline rich motifs^35^ binding to the two SH3 domains in Grb2^41^. The SAM68:Grb2 interaction may potentiate the residency time of Grb2 within the granules to activate Sos^42^ and thus drive productive Ras activity. Our finding that co-inhibition of SAM68 and Alk leads to enhanced cancer cell death and inhibition of Ras-LOCKR-S-measured Ras activity (**Fig. 4g-h**) suggests that co-treatment with Alk and SAM68 inhibitors could help overcome drug resistance in EML4-Alk-driven lung cancer^30^.

Overall, these results illustrate the power of our sensor development methodology to generate sensors for intracellular mapping of the activities, mechanisms, and functions for physiologically relevant molecules such as Ras. We envision that the tools and methods described here can be used to design a wide range of new sensors for many intracellular protein targets and thus contribute to biological and translational discovery.

## Methods

### Computational grafting of sensing domains onto latch domain

The first 7-11 amino acids from the RasBD of CRaf was grafted using Rosettascripts GraftSwitchMover into all α-helical registers between residues 325 and 359 of the latch domain within the Cage protein. The resulting Cages were energy-minimized using Rosetta fast relax, visually inspected, and typically less than ten designs were selected for subsequent cellular characterization. See **Supplementary Table 4** for details of software.

### Plasmid construction

All plasmids constructed here are using the pcDNA 3.1 backbone (unless otherwise indicated) and were produced by GenScript. See **Supplementary Table 4** for details of reagents.

### Cell culture and transfection

HEK293T, HEK293F, HEK293-FlpIn TRex, and HeLa were cultured in Dulbecco’s modified Eagle medium (DMEM) containing 1 g L^-1^ glucose and supplemented with 10% (v/v) fetal bovine serum (FBS) and 1% (v/v) penicillin–streptomycin (Pen-Strep). Beas2B, Jurkat, H3122, H2228 were cultured in Roswell Park Memorial Institute 1640 (RPMI 1640) with 10% (v/v) FBS and 1% Pen-Strep. All cells were grown in a humidified incubator at 5% CO_2_ and at 37°C.

Before transfection, all cells were plated onto sterile poly-D-lysine coated plates or dishes and grown to 50%–70% confluence. HEK293T and Beas2B cells were transfected using Turbofectin 8, HEK293F cells were transfected with PEI-MAX, Jurkat cells were transfected with Lipofectamine LTX, all other cells/conditions were transfected with Fugene HD and grown for an additional 16-24 hr before imaging. All cells underwent serum starvation for 16 hr unless indicated. See **Supplementary Table 4** for details of reagents.

### General procedures for bacterial protein production and purification

Except for purification of RBD, the *E. coli* Lemo21(DE3) strain was transformed with a pET29b^+^ plasmid encoding the synthesized gene of interest. Cells were grown for 24 hr in liquid broth medium supplemented with kanamycin. Cells were inoculated at a 1:50 ml ratio in the Studier TBM-5052 autoinduction medium supplemented with kanamycin, grown at 37°C for 2–4 hr and then grown at 18°C for an additional 18 hr. Cells were collected by centrifugation at 4,000 *g* at 4 °C for 15 min and resuspended in 30 ml lysis buffer (20 mM Tris-HCl, pH 8.0, 300 mM NaCl, 30 mM imidazole, 1 mM PMSF and 0.02 mg ml^-1^ DNase). Cell resuspensions were lysed by sonication for 2.5 min (5 s cycles). Lysates were clarified by centrifugation at 24,000 *g* at 4°C for 20 min and passed through 2-ml Ni-NTA nickel resin pre-equilibrated with wash buffer (20 mM Tris-HCl, pH 8.0, 300 mM NaCl and 30 mM imidazole). The resin was washed twice with 10 column volumes (Cversus) of wash buffer, and then eluted with 3 Cversus elution buffer (20 mM Tris-HCl, pH 8.0, 300 mM NaCl and 300 mM imidazole). The eluted proteins were concentrated using Ultra-15 Centrifugal Filter Units and further purified by using a Superdex 75 Increase 10/300 GL size exclusion column in TBS (25 mM Tris-HCl, pH 8.0, and 150 mM NaCl). Fractions containing monomeric protein were pooled, concentrated and snap-frozen in liquid nitrogen and stored at −80°C. See **Supplementary Table 4** for details of reagents.

### Procedure to purify RBD from mammalian cells

RBD proteins were produced in HEK293F cells grown in suspension using HEK293F expression medium at 33 °C, 70% humidity, 8% CO_2_ rotating at 150 rpm. The cultures were transfected using PEI-MAX with cells grown to a density of 3x10^6^ cells mL^-1^ and cultivated for 3 days. Supernatants were clarified by centrifugation (5min at 4,000 *g*), addition of polydiallyldimethylammonium chloride solution to a final concentration of 0.0375% and a second spin (5min at 4,000g).

His-tagged RBD was purified from clarified supernatants via a batch bind method, where each clarified supernatant was supplemented with 1 M Tris-HCl, pH 8.0, to a final concentration of 45 mM and 5 M NaCl to a final concentration of 310 mM. Talon cobalt affinity resin was added to the treated supernatants and allowed to incubate for 15 min with gentle shaking. Resin was collected using vacuum filtration with a 0.2-mm filter and transferred to a gravity column. The resin was washed with 20 mM Tris, pH 8.0, 300 mM NaCl, and the protein was eluted with 3 Cversus of 20 mM Tris, pH 8.0, 300 mM NaCl and 300 mM imidazole. The batch bind process was then repeated and the first and second elutions combined. SDS-PAGE was used to assess purity. Following immobilized metal affinity chromatography purification, the elution was concentrated and applied to a Cytiva S200 Increase column equilibrated with 20 mM Tris 150 mM NaCl, pH 8.0, and the peak of interest was collected and quantified using A280. See **Supplementary Table 4** for details of reagents.

### Cell counting to measure cell proliferation

Jurkat, Beas2B, H3122, and H2228 cell lines were seeded in 6-wells plates at 10,000 cells/well. Cell numbers were quantified using a hemacytometer each day for 7 days.

### Colony formation assay

Beas2B, H3122, and H2228 cell lines were seeded in 24-well plates at 100 cells/well. After 1-2 weeks to allow cell growth, cells were washed once with PBS, fixed with 4% paraformaldehyde (PFA) in PBS for 10 min, stained with 2.5 mg/mL crystal violet stain dissolved in 20% methanol for 10 min, and then washed 6x with PBS. Images were captured using the ZOE Fluorescent Cell Imager (BioRad).

### Immunostaining

293T, HeLa, Jurkat, and Beas2B cell lines were seeded onto 24-well glass-bottom plates. After transfection and drug addition, cells were fixed with 4% PFA in 2x PHEM buffer (60 mM PIPES, 50 mM HEPES, 20 mM EGTA, 4 mM MgCl_2_, 0.25 M sucrose, pH 7.3) for 10 min, permeabilized with 100% methanol for 10 min, washed with PBS 3x, blocked in 1% BSA in PBS for 30 min, incubated with primary antibody overnight at 4°C, washed with PBS 3x, incubated with DAPI, neutravidin-DyLight 650, and secondary antibody for 1 hr at room temperature and aluminum foil cover. Cells were then washed with PBS 3x and mounted for epifluorescence imaging. All images were analyzed in ImageJ. See **Supplementary Table 4** for details of reagents.

### Immunoblotting and immunoprecipitation

Cells expressing indicated constructs and incubated with indicated drugs were plated, transfected, and labeled as described in figure legends. Cells were then transferred to ice and washed 2x with ice cold DPBS. Cells were then detached from the well by addition of 1x RIPA lysis buffer (50 mM Tris pH 8, 150 mM NaCl, 0.1% SDS, 0.5% sodium deoxycholate, 1% Triton X-100, 1x protease inhibitor cocktail, 1 mM PMSF, 1mM Na_3_VO_4_, 1% NP-40) and either scraping of cells or rotation on shaker for 30 min at 4°C. Cells were then collected and vortexed for at least 5 s every 10 min for 20 min at 4°C. Cells were then collected and clarified by centrifugation at 20,000 rpm for 10 minutes at 4°C. The supernatant was collected and underwent Pierce BCA assay to quantify total protein amounts.

For immunoblotting, whole cell lysate protein amounts were normalized across samples in the same gel, mixed with 4x loading buffer prior to loading, incubated at 95°C for 5 min and then 4°C for 5 min, and separated on Any kDa SDS-PAGE gels. Proteins separated on SDS-page gels were transferred to nitrocellulose membranes via the TransBlot system (BioRad). The blots were then blocked in 5% milk (w/v) in TBST (Tris-buffered saline, 0.1% Tween 20) for 1 hr at room temperature. Blots were washed with TBST 3x then incubated with indicated primary antibodies in 1% BSA (w/v) in TBST overnight at 4°C. Blots were then washed with TBST 3x and incubated with LICOR dye-conjugated secondary antibodies (LICOR 680/800 or streptavidin-LICOR 800) in 1% BSA (w/v) in TBST for 1 hr at room temperature. The blots were washed with TBST 3x and imaged on an Odyssey IR imager (LICOR). Quantitation of Western blots was performed using ImageJ on raw images.

For immunoprecipitation, agarose beads were either preloaded with streptavidin (high capacity streptavidin beads) or loaded by 3x lysis buffer washes and then addition of 1mg ml^-1^ indicated antibodies at 4°C on orbital shaker for 3 hr. Beads were then washed 2x in lysis buffer. Whole cell lysate protein amounts were normalized across samples and protein samples were added to beads (at least 100µg per sample) either at room temperature for 1 hr for streptavidin beads or at 4°C on orbital shaker overnight. Beads were then washed 2x in lysis buffer and 1x in TBS and then mixed with 4x loading buffer sometimes containing 2mM biotin and 20mM DTT^43^ for streptavidin pulldowns. The remaining portion of the protocol is the same as immunoblotting. See **Supplementary Table 4** for details of reagents.

### Mass spectrometry analysis

Cells expressing indicated constructs and incubated with indicated drugs were plated, transfected, and labeled as described in figure legends. Cells were then transferred to ice and washed 2x with ice cold DPBS, detached from the well by addition of 1x RIPA lysis buffer (50 mM Tris pH 8, 150 mM NaCl, 0.1% SDS, 0.5% sodium deoxycholate, 1% Triton X-100, 1x protease inhibitor cocktail, 1 mM PMSF, 1mM Na_3_VO_4_, 1% NP-40) and scraping of cells, collected and vortexed for at least 5 s every 10 min for 20 min at 4°C, and collected and clarified by centrifugation at 20,000g for 10 minutes at 4°C. The supernatant was collected and underwent Pierce BCA assay to quantify total protein amounts.

50µL of high capacity streptavidin agarose beads were washed 2x in lysis buffer. Whole cell lysate protein amounts were normalized across samples and protein samples were added to beads (at least 100µg per sample) at room temperature for 1 hr. Beads were then washed 2x with lysis buffer, 1x with 1M KCl, 1x with 0.1M Na_2_CO_3_, 2x 2M urea, and 2x with TBS. Beads were re-suspended in 50µL of denaturing buffer (6M guanidinium chloride, 50mM Tris containing 5mM TCEP and 10mM CAM with TCEP and CAM added fresh every time), inverted a few times, and heated to 95°C for 5 min. The bead slurry was diluted with 50µL of 100mM TEAB and 0.8µg of LysC was added per sample with the pH adjusted to 8-9 using 1M NaOH. This mixture was agitated on a thermomixer at 37°C for 2 hr at 1400 rpm. Afterwards, samples were diluted 2x with 100µL of 100mM TEAB with 0.8µg of sequencing grade trypsin per sample with the pH adjusted to 8-9 using 1M NaOH. This mixture was agitated on a thermomixer at 37°C for 12-14 hr at 800 rpm. After overnight trypsinization, samples were diluted 2x with 200µL of Buffer A (5% acetonitrile with 0.1% TFA) containing 1% formic acid. These samples were inverted a few times and pH adjusted to 2-3 using 100% formic acid. StageTips for peptide desalting were prepared by extracting out plugs from C18 matrices, shoved down a 200µL tip, and pressed with a plunger for flatness. Using these StageTips, 50µL of Buffer B (80% acetonitrile with 0.1% TFA) was passed through at 4000g for 1 min followed by 5µ0L of Buffer A for 4000g for 1 min. The supernatant of the samples was added to StageTips and spun down at 4000g for 5 min. Then, 50µL of Buffer A was added and spun down at 4000g for 2.5 min. 50µL of Buffer B was added to stage tips and a syringe pump was applied to elute samples.

Peptide samples were separated on an EASY-nLC 1200 System (Thermo Fisher Scientific) using 20 cm long fused silica capillary columns (100 µm ID, laser pulled in-house with Sutter P-2000, Novato CA) packed with 3 μm 120 Å reversed phase C18 beads (Dr. Maisch, Ammerbuch, DE). The LC gradient was 90 min long with 5−35% B at 300 nL/min. LC solvent A was 0.1% (v/v) aqueous acetic acid and LC solvent B was 20% 0.1% (v/v) acetic acid, 80% acetonitrile. MS data was collected with a Thermo Fisher Scientific Orbitrap Fusion Lumos using a data-dependent data acquisition method with a Orbitrap MS1 survey scan (R=60K) and as many Orbitrap HCD MS2 scans (R=30K) possible within the 2 second cycle time.

### Computation of MS raw files

Data .raw files were analyzed by MaxQuant/Andromeda version 1.5.2.8 using protein, peptide and site FDRs of 0.01 and a score minimum of 40 for modified peptides, 0 for unmodified peptides; delta score minimum of 17 for modified peptides, 0 for unmodified peptides. MS/MS spectra were searched against the UniProt human database (updated July 22nd, 2015). MaxQuant search parameters: Variable modifications included Oxidation (M) and Phospho (S/T/Y). Carbamidomethyl (C) was a fixed modification. Max. missed cleavages was 2, enzyme was Trypsin/P and max. charge was 7. The MaxQuant “match between runs” feature was enabled. The initial search tolerance for FTMS scans was 20 ppm and 0.5 Da for ITMS MS/MS scans.

### MaxQuant output data processing

MaxQuant output files were processed, statistically analyzed and clustered using the Perseus software package v1.5.6.0. Human gene ontology (GO) terms (GOBP, GOCC and GOMF) were loaded from the ‘mainAnnot.homo_sapiens.txt’ file downloaded on 02.03.2020. Expression columns (protein and phosphopeptide intensities) were log2 transformed and normalized by subtracting the median log2 expression value from each expression value of the corresponding data column. Potential contaminants, reverse hits and proteins only identified by site (biotinylation) were removed. Reproducibility between LC-MS/MS experiments was analyzed by column correlation (Pearson’s r) and replicates with a variation of r > 0.25 compared to the mean r-values of all replicates of the same experiment (cell line or knockdown experiment) were considered outliers and excluded from the analyses. Data imputation was performed in Perseus using a modeled distribution of MS intensity values downshifted by 1.8 and having a width of 0.2. Hits were further filtered using gene ontology analysis (signaling pathways) via Panther database. See **Supplementary Table 4** for details of reagents.

### *In vitro* fluorescence characterization

A Synergy Neo2 Microplate Reader (BioTek) was used for all in vitro fluorescence measurements. Assays were performed in 1x PBS. The purified protein components (+50µM DFHBI-1T for mFAP2a experiments) were placed in 96-well black-well clear-bottom plates, centrifuged at 1,000g for 1 min, and incubated for 30 min at room temperature to enable pre-equilibration. Fluorescence measurements in the absence of target were taken every 1 min after injection (0.1 s integration and 10 s shaking during intervals) at the indicated wavelengths: For FRET spectra, the wells were excited at wavelengths indicated in figure legends and the respective FRET was recorded at 5 nm intervals. See **Supplementary Table 4** for details of reagents.

### Time-lapse epifluorescence imaging

Cells were washed twice with FluoroBrite DMEM imaging media and subsequently imaged in the same media in the dark at room temperature. Forskolin, EGF, A115, a- CD3E + a-CD28 were added as indicated. Epifluorescence imaging was performed on a Yokogawa CSU-X1 spinning dish confocal microscope with either a Lumencor Celesta light engine with 7 laser lines (408, 445, 473, 518, 545, 635, 750 nm) or a Nikon LUN-F XL laser launch with 4 solid state lasers (405, 488, 561, 640 nm), 40x/0.95 NA objective and a Hamamatsu ORCA-Fusion scientific CMOS camera, both controlled by NIS Elements 5.30 software (Nikon). The following excitation/FRET filter combinations (center/bandwidth in nm) were used: CFP: EX445 EM483/32, CFP/YFP FRET: EX445 EM542/27, YFP: EX473 EM544/24, GFP: EX473 EM525/36, RFP: EX545 EM605/52, Far Red (e.g. AlexaFluor 647): EX635 EM705/72, 450: EX445 EM525/36, 500: EX488 EM525/36. Exposure times were 100 ms for acceptor direct channel and 500ms for all other channels, with no EM gain set and no ND filter added. Cells that were too bright (acceptor channel intensity is 3 standard deviations above mean intensity across experiments) or with significant photobleaching prior to drug addition were excluded from analysis. All epifluorescence experiments were subsequently analyzed using Image J. Brightfield images were acquired on the ZOE Fluorescent Cell Imager (BioRad). See **Supplementary Table 4** for details of reagents.

### FRET biosensor analysis

Raw fluorescence images were corrected by subtracting the background fluorescence intensity of a cell-free region from the FRET intensities of biosensor-expressing cells. Cyan/yellow FRET ratios were then calculated at each time point (*R)*. For some curves, the resulting time courses were normalized by dividing the FRET ratio at each time point by the basal ratio value at time zero (*R*/*R*_0_), which was defined as the FRET ratio at the time point immediately preceding drug addition (*R*_0_)^44^. Graphs were plotted using GraphPad Prism 8 (GraphPad).

### Co-localization analysis

For co-localization analysis, cell images were individually thresholded and underwent Coloc 2 analysis on ImageJ. Mander’s coefficient, which ranges from 0 to 1 with 1 being 100% colocalized, is measuring the spatial overlap of one imaging channel (e.g. EML4-Alk-YFP) with another imaging channel (e.g. immunostained SAM68). Pearson’s coefficient compares the pixel intensity of one channel with another channel. Pearson’s coefficient values can range from -1 to 1 with -1 meaning inversely proportional and 1 meaning same pixel intensities.

### Quantification of cellular puncta

For analysis of puncta number, cell images were individually thresholded and underwent particle analysis with circularity and size cutoffs in ImageJ.

### Statistics and reproducibility

No statistical methods were used to predetermine the sample size. No sample was excluded from data analysis, and no blinding was used. All data were assessed for normality. For normally distributed data, pairwise comparisons were performed using unpaired two-tailed Student’s t tests, with Welch’s correction for unequal variances used as indicated. Comparisons between three or more groups were performed using ordinary one-way or two-way analysis of variance (ANOVA) as indicated. For data that were not normally distributed, pairwise comparisons were performed using the Mann-Whitney U test, and comparisons between multiple groups were performed using the Kruskal-Wallis test. All data shown are reported as mean ± SEM and error bars in figures represent SEM of biological triplicates. All data were analyzed and plotted using GraphPad Prism 8 including non-linear regression fitting.

### Data availability

The data that support the findings of this study are available from the corresponding author upon reasonable request. All accession codes have been provided for the paper. Source data are provided with this paper.

### Code availability

Source data are provided with this paper.

## Acknowledgements

We acknowledge funding from HHMI (J.Z.Z. and D.B.), Helen Hay Whitney Foundation (J.Z.Z.), the Audacious Project at the Institute for Protein Design (J.Z.Z, D.B.), NIH grants (R01GM129090 (S-E.O.), R01GM145011 (D.J.M.), and R01GM086858 (D.J.M.)). This work used an EASY-nLC1200 UHPLC and Thermo Scientific Orbitrap Fusion Lumos Tribrid mass spectrometer purchased with funding from a National Institutes of Health SIG grant S10OD021502. We thank J.C. Klima, R.A. Langan, and S.E. Boyken for discussion on LOCKR sensor development, B. Fiala at the Institute for Protein Design for providing SARS-CoV-2 RBD and LCB1, A. Luis for help in processing mass spectrometry samples, I. C. Haydon for providing some schematics in the manuscript, A.Y. Ting for discussion on the split proximity labeler, R. Bayliss and J. Sampson for EML4-Alk constructs and cell lines, M. Ahlrichs for help with mammalian cells and cell culture, and M.R. Philips and T.G. Bivona for fruitful discussion of the Ras results.

## Author contributions

J.Z.Z. conceived of the sensor and project. J.Z.Z., D.J.M., and D.B. supervised, designed, and interpreted the experiments. J.Z.Z. and W.H.N. performed all experiments. N.G. helped with structure predictions. J.C.R. made CIAR-Golgi cell lines. S.E.O. ran samples through mass spectrometry. J.Z.Z., D.J.M, and D.B. wrote the original draft. All authors reviewed and commented on the manuscript.

## Competing interest

J.Z.Z., D.J.M., and D.B. are co-inventors in a provisional patent application (application number 63/380,884 submitted by the University of Washington) covering the biosensors described in this manuscript.

## Extended Data Figures

**Extended Data Figure 1:**
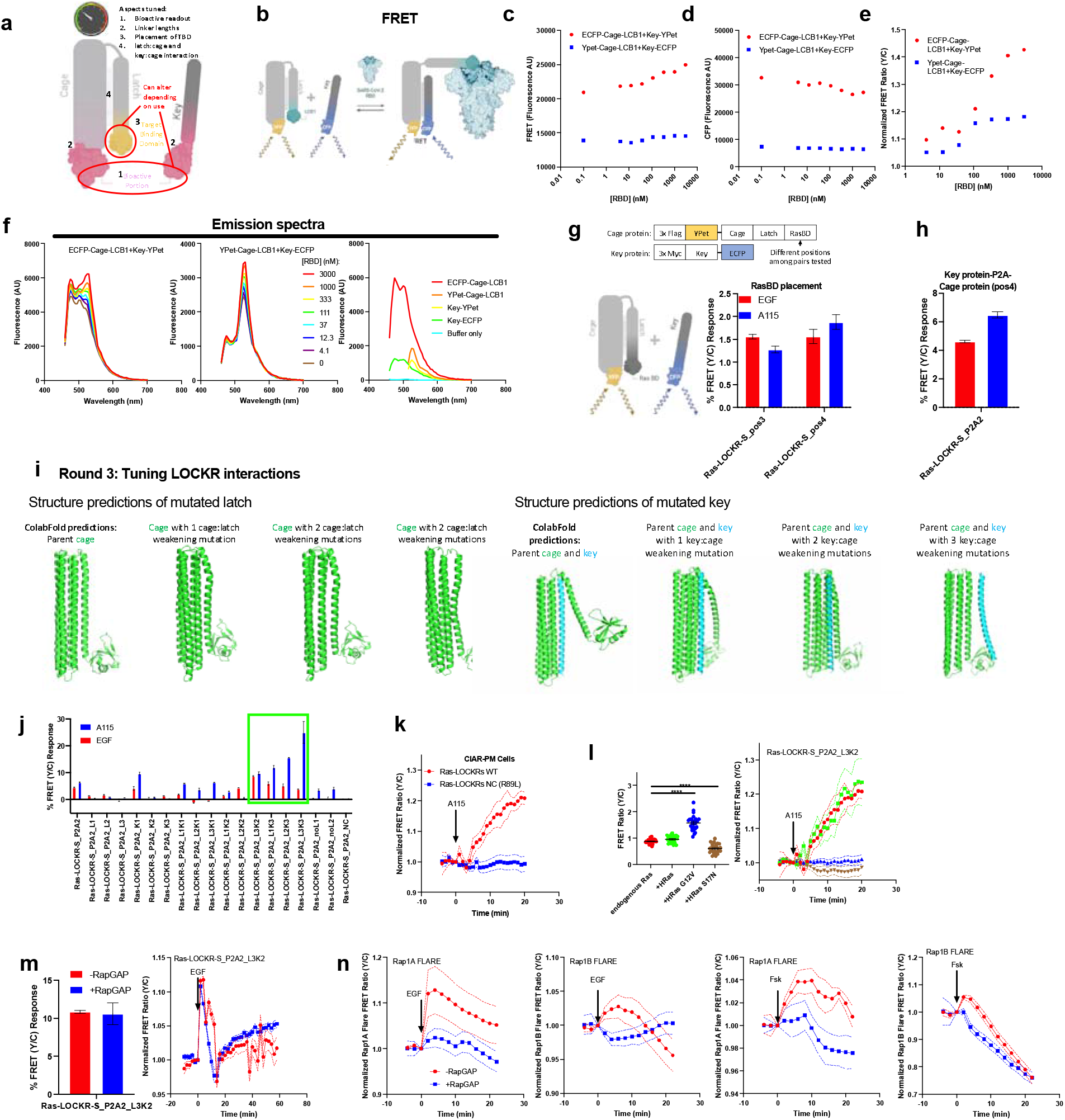
Design and characterization of Ras-LOCKR-S to track compartmentalized endogenous Ras activity dynamically and specifically. **a**, Schematic describing the different aspects of Ras-LOCKR-S that were tuned. **b-e,** FRET-based LOCKR sensors modifying lucCageRBD, which senses RBD of SARS-CoV-2 spike protein via binding to the *de novo* protein binder LCB1 (Cao et al., 2020), (**b**) with CFP and YFP as FRET donor and acceptor, respectively. Two placements of the FPs were tested *in vitro* for yellow/cyan FRET fluorescence (**c**), CFP fluorescence (**d**), and normalized yellow/cyan FRET ratio changes (normalized to no RBD) (**e**) with a range of RBD concentrations ([Cage]=[Key]=1mM). **f**, FRET spectra (excitation wavelength: ECFP+YPet, ECFP only, buffer only=450nm, YPet only=510nm) of the two Key-Cage pairs against a range of RBD concentrations ([Cage]=[Key]=1mM) (left and middle) and the Key or Cage proteins alone (right). **g**, (top) Domain structures shown. (bottom) Various FRET-based Ras-LOCKR-S candidates with different RasBD grafting positions on latch using Rosetta-based GraftSwitchMover (see Methods) were tested in 293T cells (100ng/mL EGF stimulation) and CIAR-PM (250nM A115 stimulation) cells with percent FRET ratio changes reported (n=at least 21 cells per condition). Only YFP-tagged Cage and CFP-tagged Key were tested as YFP-tagged Key did not show fluorescence. **h**, RasBD placements identified in Round 1 were used in unimolecular (only Cage, no Key) without linker (uni), with linker (uniL), and bimolecular (with P2A sequence) formats. One of the bimolecular designs showed consistent responses to EGF and A115 (n=at least 18 cells per condition). **i**, AlphaFold structure predictions of Ras-LOCKR-S Cage only (left) or Key:Cage complex (right) with the indicated mutations. **j**, Point mutations to weaken latch:cage or key:cage interactions (the N-terminal half of latch and key are similar) are identified via structure predictions and implemented in the bimolecular Ras-LOCKR-S candidate identified in **h**. The Ras-LOCKR-S candidates that display consistent FRET ratio changes to EGF and A115 are boxed in green and were further validated in subsequent experiments. **k**, Negative controls (NC, RasBD^R89L^) shows negligible FRET ratio changes (bar: n=at least 19 cells per condition, time-course: n=at least 10 cells per condition). **l**, CIAR-PM-239T cells expressing Ras-LOCKR-S_P2A2_L3K2 and co-expressing either HRas, constitutively active HRas (HRas^G12V^), or dominant negative HRas (HRas^S17N^) are stimulated with 250 nM A115. FRET ratio changes (left, n=at least 11 cells per condition) and starting FRET ratios (right, n=25 cells per condition) are reported. **m**, Ras-LOCKR-S_P2A2_L3K2 was tested for Ras but not Rap selectivity in 293T cells expressing RapGAP and stimulated with EGF (left, n=at least 25 cells per condition) and with expected Ras dynamics (right, n=13 cells per condition). **n**, Testing of RapGAP^45^ in 293T cells using Rap1A and Rap1B FLARE activity reporters^19^. FRET ratio changes of 293T cells expressing either Rap1A FLARE or Rap1B FLARE with and without RapGAP co-expression are stimulated with either EGF (left) or 50µM Forskolin (Fsk) (n=at least 25 cells per condition). See **Supplementary Table 1** for domain structures and sequences. For all graphs, solid lines indicate representative average time courses of FRET ratio changes with error bars representing standard error mean (SEM). Bar graphs represent mean + SEM. ****p < 0.0001, ordinary one-way ANOVA.

**Extended Data Figure 2:**
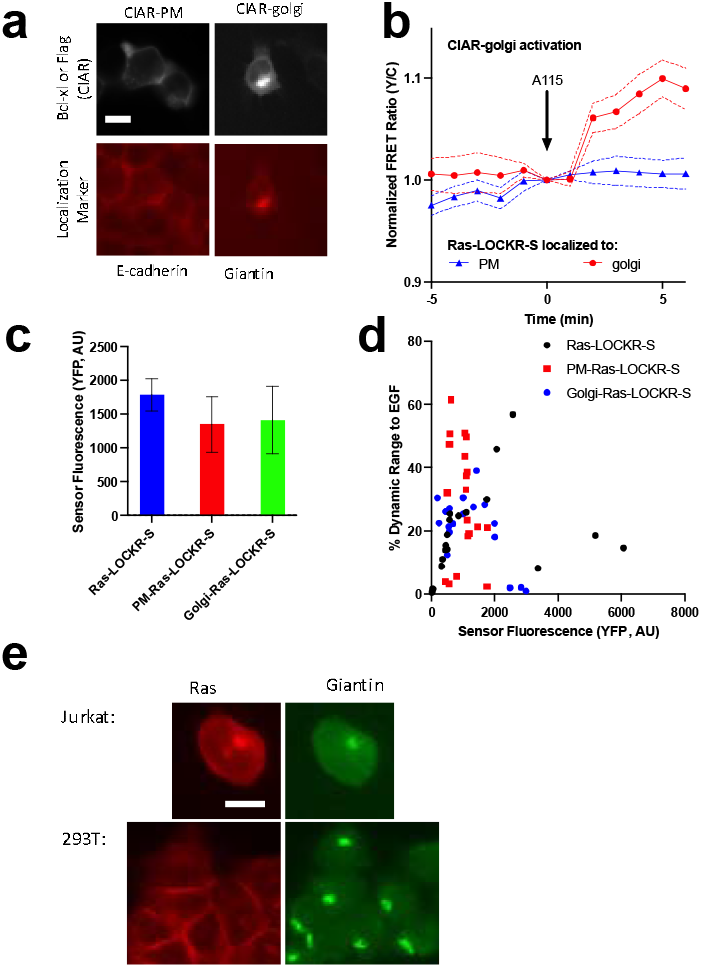
Characterization of localized Ras-LOCKR-S and CIAR in 293T and Jurkat cells. **a**, CIAR localized to PM (KRas4a CAAX) or Golgi (Giantin^3131-3259^)^46^ is stably expressed in 293-TRex cells, which were immunostained for CIAR via Bcl-xL (CIAR-PM) or Flag tag (CIAR-Golgi) and their respective localization markers with representative epifluorescence images shown. **b**, FRET ratio changes after A115 addition onto cells stably expressing Golgi-localized CIAR (CIAR-Golgi) and transfected with subcellularly localized Ras-LOCKR-S (n=9 cells per condition). **c**, Average localized Ras-LOCKR-S (Cage, YFP) fluorescence. **d**, Scatterplot of sensor fluorescence (Cage, YFP) compared to % dynamic range to EGF. **e**, Representative epifluorescence images of Jurkat T-cells and 293T cells immunostained for Ras and giantin (Golgi marker). Solid lines indicate representative average time courses of FRET ratio changes with error bars representing standard error mean (SEM). Bar graphs represent mean ± SEM. Scale bars = 10µm.

**Extended Data Figure 3:**
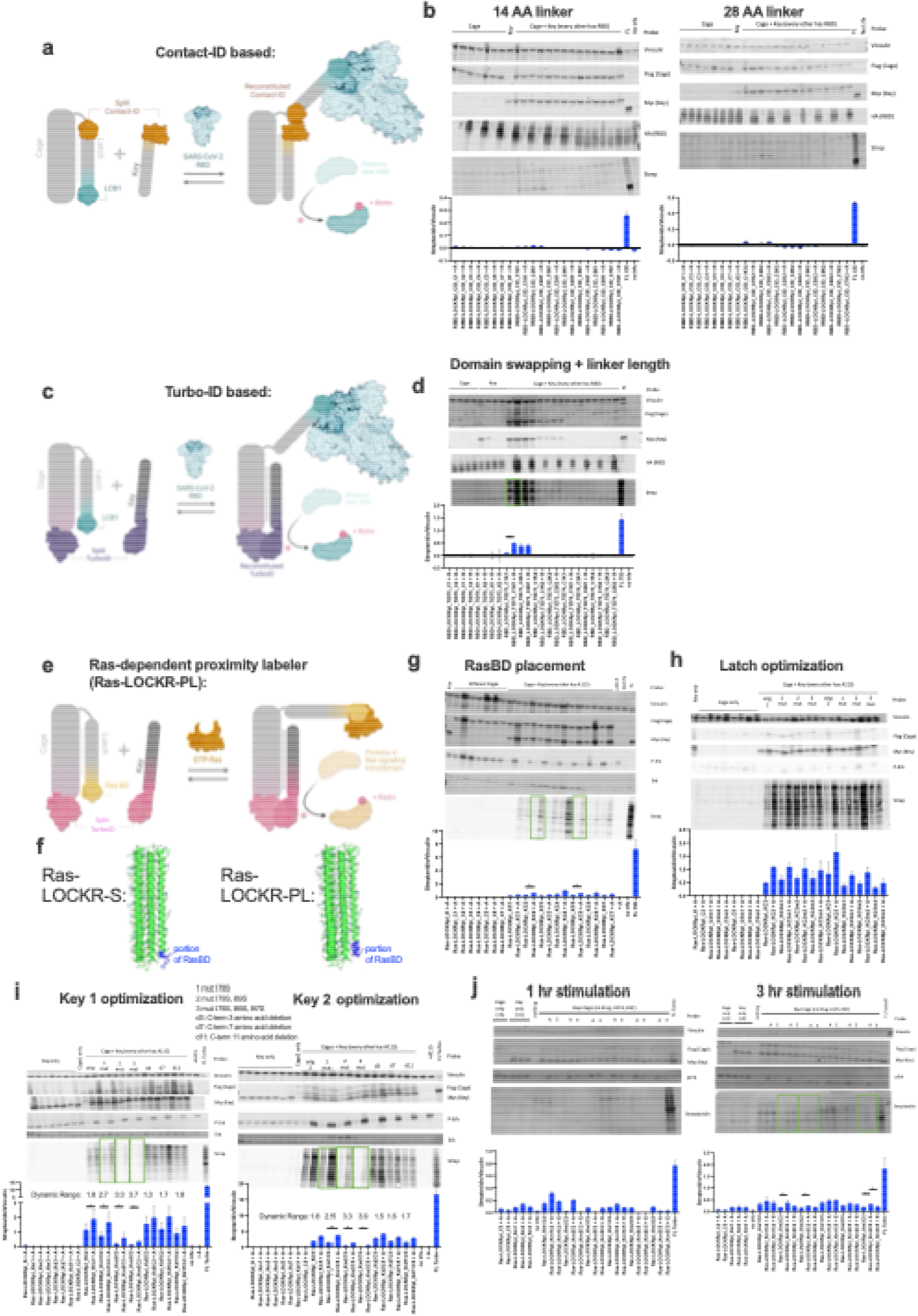
Development of Ras-LOCKR-PL. **a-b**, Split ContactID-based LOCKR proximity labelers were tested using similar architecture to lucCageRBD. Schematic in **a** and data shown in **b**. Different grafting positions of the smaller bit of ContactID on the latch using Rosetta-based GraftSwitchMover (see Methods) and different linker lengths between larger bit of ContactID and Key were tested in 293T cells transfected with Flag-tagged Cage, Myc-tagged Key, full length ContactID (FL CID), and HA-tagged RBD (receptor binding domain of SARS-CoV-2 spike protein) with 5x nuclear exclusion sequence. After 16 hr 500µM biotin incubation, these cells were subsequently immunoblotted for the transfected proteins and biotinylation via streptavidin. **c-d**, Split TurboID (split site TurboID^73/74^)-based LOCKR proximity labelers modifying lucCageRBD were tested in 293T cells transfected with Cage, Key, full length TurboID (FL TID), and/or RBD and immunoblotted for biotinylation. Different placements of the split TurboID and linker lengths between split TurboID and Cage/Key were tested with the Ras-LOCKR-PL candidate that led to the highest increase in biotinylation upon RBD expression boxed in green and optimized further. Schematic in **c** and data shown in **d**. **e-j**, Testing of Ras-LOCKR-PL candidates was done via western blotting of CIAR-PM-239T cells incubated with 250nM A115 (labeled A) or 100ng/mL EGF (labeled E) and 500µM biotin for either 16 hr (**e-i**), 1 hr (**j**, left), or 3 hr (**j**, right). RasBD was grafted onto the latch using GraftSwitchMover (**f**, shows structure predictions) and tested for increases in biotinylation after Ras activation by A115 (FL = full length TurboID). The highest dynamic range Ras-LOCKR-PL candidates are boxed in green (**g**) and further optimized by either weakening latch:cage interaction by mutating latch (**h**) or weakening key:cage interaction by mutating the Key (2 different Cages tested for left and right) (**i**). C-terminal Key truncations were also tested to weaken key:cage interaction (**i**). The highest dynamic range Ras-LOCKR-PL candidates boxed in green (**i**) were tested in shorter time regimes either with A115 or EGF (**j**). See **Supplementary Table 2** for domain structures and sequences. All immunoblots are representative of at least 3 experiments.

**Extended Data Figure 4:**
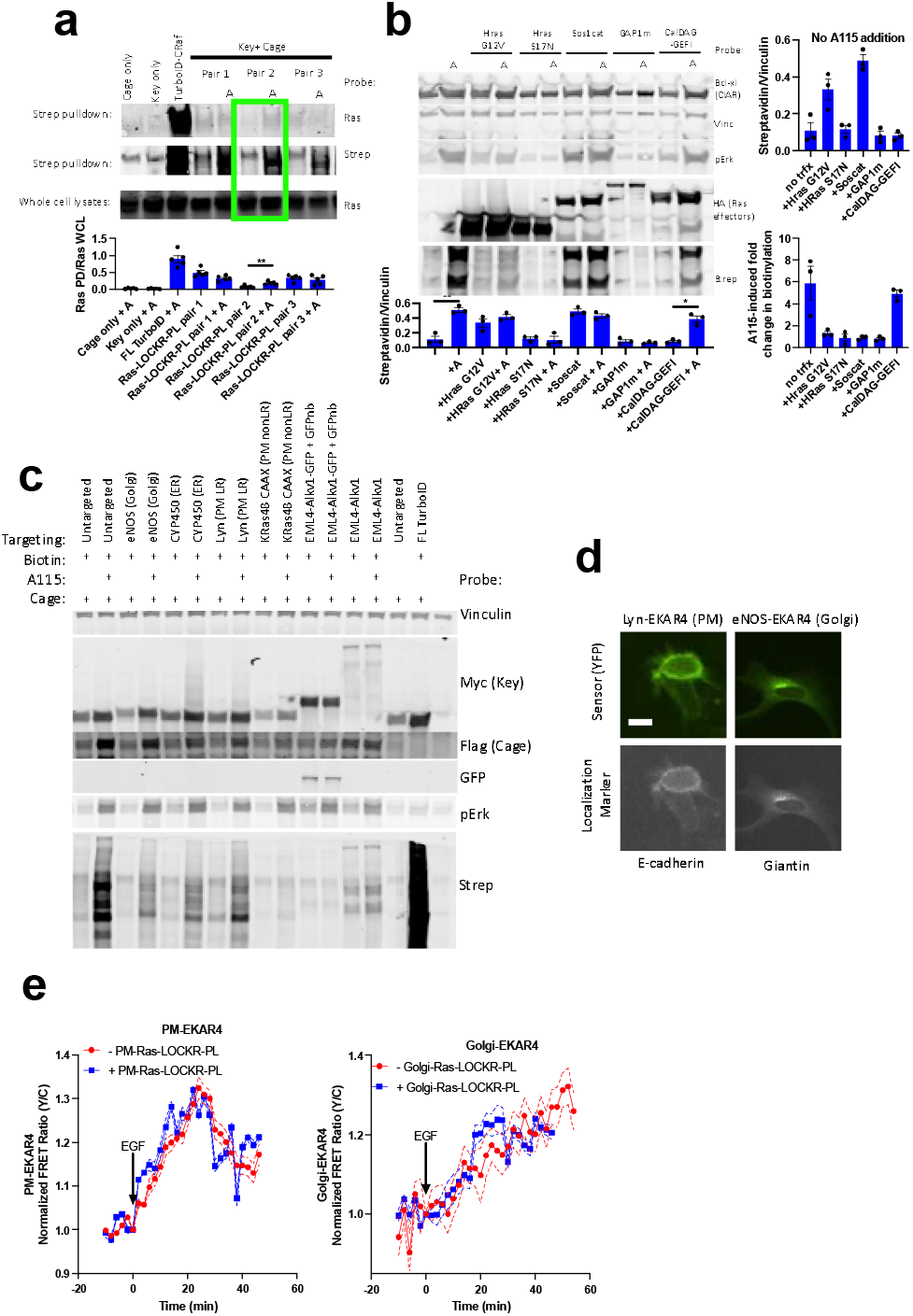
Characterization of Ras-LOCKR-PL. **a-c**, Representative immunoblots of CIAR-PM cells with 500µM biotin with or without 250nM A115 (labeled “A”) for 16 hours. (**a**) Negative controls (Myc-tagged Key or Flag-tagged Cage only), positive control (TurboID-CRaf), and Ras-LOCKR-PL tested designs (Key+Cage) were expressed, and treated with DMSO or with A115. Cells underwent streptavidin pulldown (PD) and probed for Ras with Ras PD over whole cell lysate (WCL) quantified (n= at least 4 experimental repeats). (**b**) Optimized Ras-LOCKR-PL co-expressed with various HA-tagged Ras mutants/effectors or Rap effector (CalDAG-GEFI). (**c**) Localized Ras-LOCKR-PLs were tested for A115-induced biotinylation in CIAR-PM-293T cells. **d**, Representative epifluorescence images of EKAR4^29^ localized to PM (N-terminus of Lyn)^22^ or Golgi (N-terminus of eNOS)^23^ transfected into 293T cells, which were immunostained for their respective localization markers. **e**, 293T cells transfected with localized EKAR4 and either with or without Ras-LOCKR-PL also localized to same region. Normalized FRET ratio changes over time of these cells stimulated with 100ng/mL EGF. Solid lines indicate representative average time with error bars representing standard error mean (SEM). Bar graphs represent mean ± SEM, n=at least 3 experiments per condition. **p < 0.01, *p < 0.05, unpaired two-tailed Student’s t-test. Scale bar = 10µm.

**Extended Data Figure 5:**
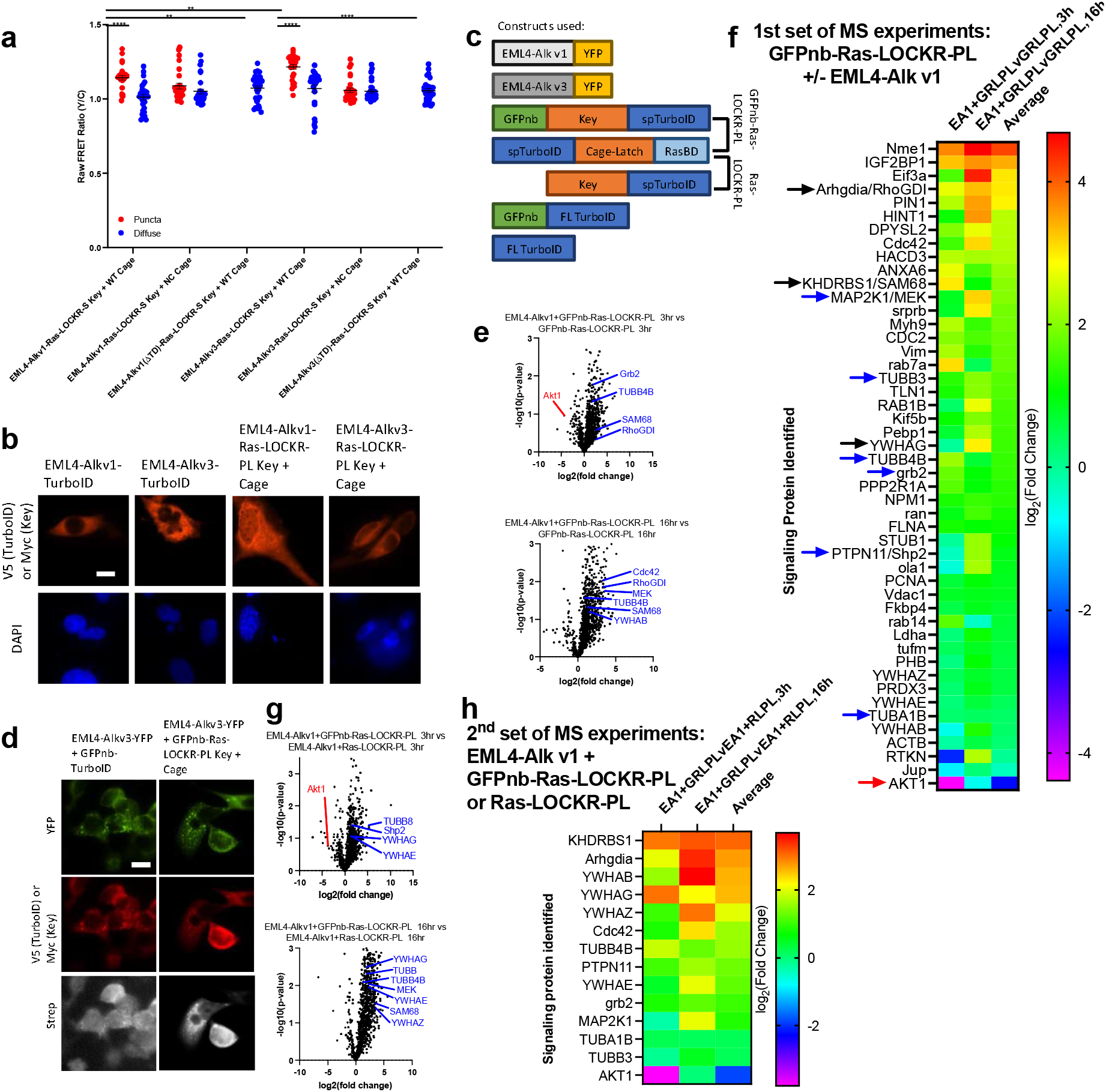
Ras-LOCKR tools identify the Ras activity and environment inside EML4-Alk granules. **a**, Raw FRET ratios of Beas2B cells transfected with Ras-LOCKR-S or NC sensor tethered to EML4-Alk variant 1 (v1)/variant 3 (v3) full length or with trimerization domain deleted (ΔTD). **b**, Representative epifluorescence images of Beas2B cells transfected with V5-tagged TurboID or Myc-tagged Ras-LOCKR-PL tethered to EML4-Alk v1/v3 and immunostained for respective tags and biotinylation via streptavidin. **c**, Domain structures of constructs used. **d**, Representative epifluorescence images of Beas2B (WT lung) transfected with EML4-Alk variant 3 (v3), full length V5-tagged TurboID tethered to GFP nanobody (GFPnb), or Myc-tagged Ras-LOCKR-PL tethered to GFPnb. **e**, Volcano plot of mass spectrometry results of Beas2B cells expressing indicated constructs and stimulated for indicated durations with 500µM biotin. Plotted differences are comparing the first condition listed in the title (right) against the second condition listed (left). **f**, From MS data, hits were filtered based on selectively labeling during EML4-Alk v1 expression (more than 2-fold change (-log_2_(student’s t-test difference) > 1)), passed a statistical cutoff (p-value cutoff of 0.5 (-log_10_p-value ∼ 1.3)), are signaling proteins as identified in gene ontology analysis, and abundant proteins (proteins related to the proteasome and ribosome) were excluded. Heat maps (colored by -log_2_student’s t-test difference) display the signaling-related proteins detected in Beas2B cells transfected with Ras-LOCKR-PL or TurboID constructs localizing to EML4-Alk v1/v3 granules and incubated with 500µM biotin for indicated times. With experimental comparisons listed on top, student’s t-test difference indicates propensity of protein to be more enriched in the first listed experimental sample compared to second with 0 indicating no difference. Blue arrows are proteins expected to be enriched in EML4-Alk v1 granules, red arrows are proteins expected to be excluded from EML4-Alk v1 granules, black arrows are new hits that were verified later on to be sequestered in EML4-Alk granules, and gray arrows are known interactors (based on STRING databases) of the validated proteins. **g**, Volcano plot of mass spectrometry results of Beas2B cells expressing indicated constructs and stimulated for indicated durations with 500µM biotin. Plotted differences are comparing the first condition listed in the title (right) against the second condition listed (left). **h**, Hits identified in first set of MS experiments (**e-f**) were identified in the second set of MS experiments (**g**), and their -log_2_(student’s t-test difference) from the second set of MS experiments are listed and colored in a heatmap. MS data is documented in **Supplementary Table 3**. Bar graphs represent mean + SEM. ****p < 0.0001, **p < 0.01, ordinary two-way ANOVA. Scale bars = 10µm.

**Extended Data Figure 6:**
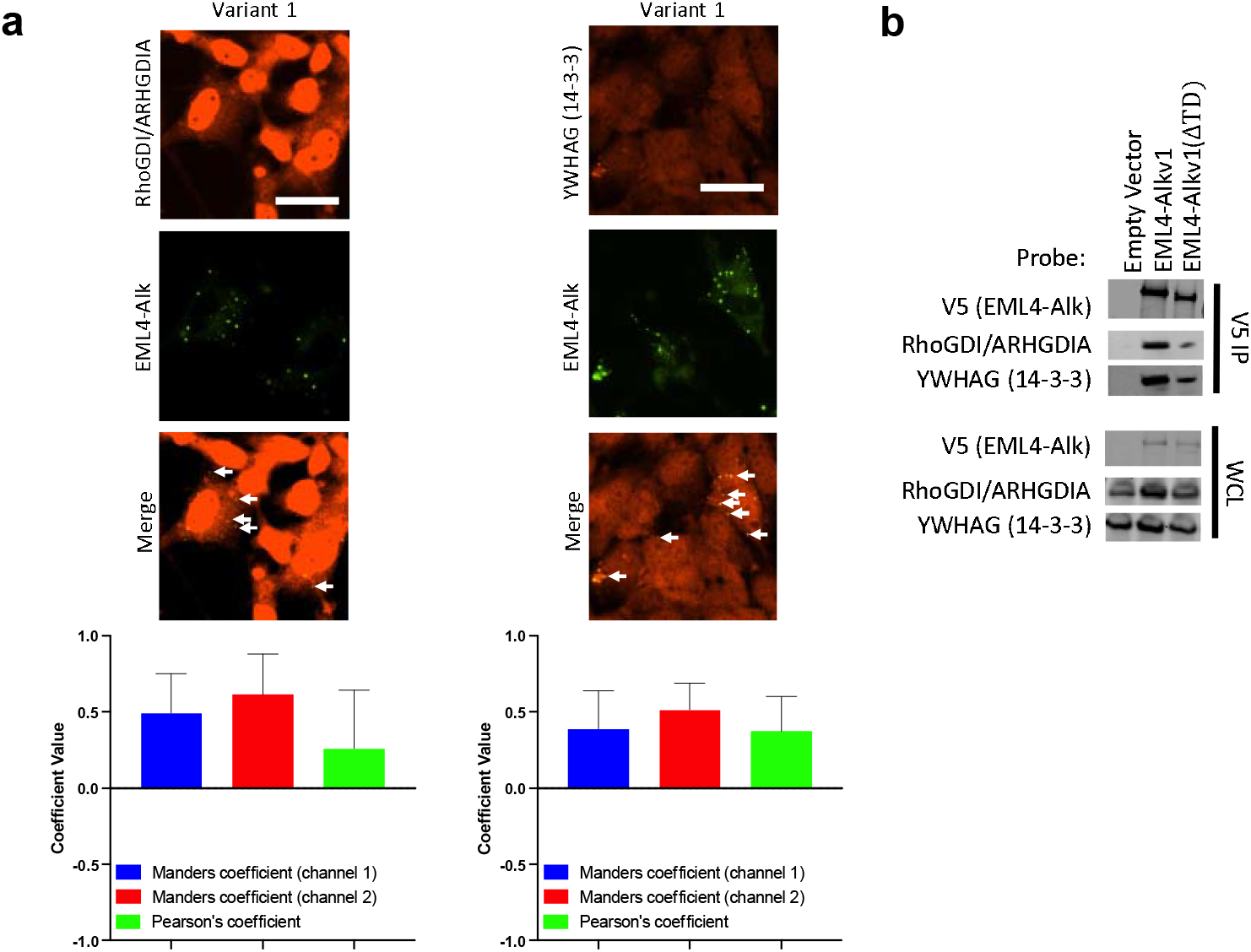
Identification of components sequestered in EML4-Alk granules. **a**, (top) Representative epifluorescence images of Beas2B cells expressing GFP-tagged EML4-Alk v1 and immunostained for hits identified in the mass spectrometry analysis. Arrows indicate co-localization of EML4-Alk with probed protein. (bottom) Colocalization analysis. **b**, Representative immunoblot of Beas2B cells transfected with V5-tagged EML4-Alk v1 full length or with trimerization domain deleted (ΔTD), subjected to V5 immunoprecipitation, and probed for hits identified in the MS analysis. Bar graphs represent mean ± SEM.

**Extended Data Figure 7:**
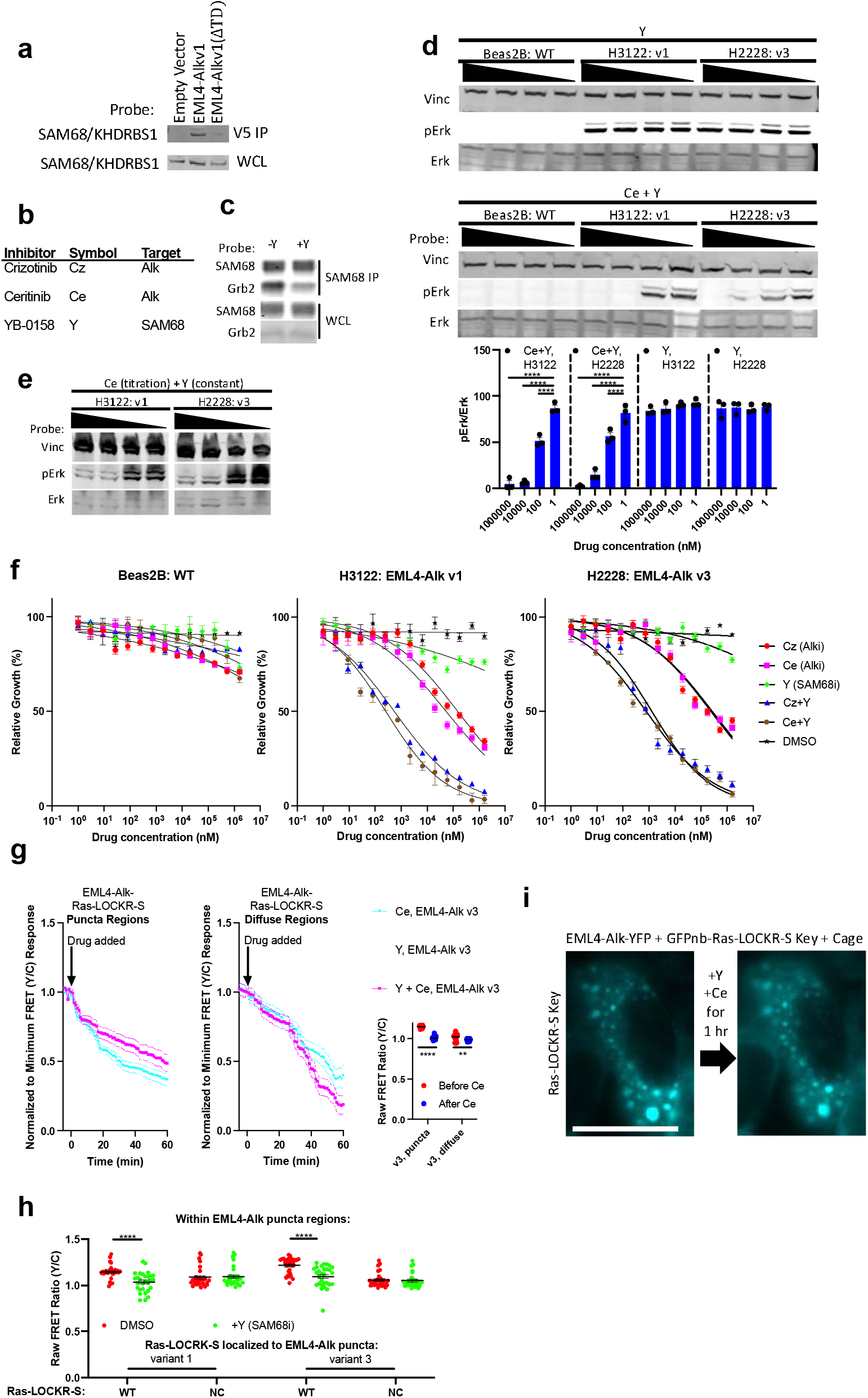
SAM68 regulates Ras signaling inside EML4-Alk granules. **a**, Representative immunoblot of Beas2B cells transfected with V5-tagged EML4-Alk v1 full length or with trimerization domain deleted (ΔTD), subjected to V5 immunoprecipitation, and probed for SAM68. **b**, List of inhibitors used, their abbreviations, and their target. **c**, Representative immunoblot of Beas2B cells treated with 1M YB-0158, underwent Sam68 immunoprecipitation, and probed for Sam68 and Grb2. Top: Sam68 immunoprecipitation. Bottom: whole cell lysate. **d**, (top) Representative immunoblot of Beas2B, H3122, and H2228 cells treated for 1 hr with varying concentration of Ceritinib (1nM-1mM) and YB-0158 (1nM-1mM). (bottom) Relative phosphorylated Erk over total Erk quantified (n=3 experiments per condition). **e**, Representative immunoblot of H3122 and H2228 cells treated for 1 hr with either constant concentration of YB-0158 (200nM) and a range of concentrations of Ceritinib (1nM-1mM) (left) or a constant concentration of Ceritinib (200nM) and a range of concentrations of YB-0158 (1nM-1mM) (right). **f**, Cell count of Beas2B, H3122, H2228 cell lines incubated for 1 week with varying concentrations of inhibitors to Alk and/or SAM68 (n=3 experiments per condition), or DMSO. **g**, (left) Normalized to minimum FRET ratio time-courses of Beas2B cells transfected with Ras-LOCKR-S localized to EML4-Alk v3 and incubated with 10µM of inhibitors. Puncta and diffuse regions were analyzed separately. Normalized to minimum FRET ratios are calculated by normalizing the data set to the condition with the largest decrease in FRET ratios (Ce + Y in both cases) where 0 represents the lowest FRET ratio out of the entire data set. (right) Raw FRET ratios of puncta and diffuse EML4-Alk regions after Alk inhibition for 1 hour. **h**, Average raw FRET ratios in the punctate regions of Beas2B cells expressing Ras-LOCKR-S WT or NC sensor fused to EML4-Alk v1. Cells were treated with either DMSO or 1µM YB-0158 for 1 hr (n=at least 15 cells per condition). **i**, Representative epifluorescence images in CFP channel (for Ras-LOCKR-S Key) of Beas2B cells expressing YFP-tagged EML4-Alk v1, GFPnb-Ras-LOCKR-S Key, and Cage. These cells were incubated with 10µM YB-0158 and Ceritinib for 1 hour. Solid lines in **f** indicate IC_50_ fit with points representing average normalized cell count. Solid lines in **g** indicate representative average time with error bars representing standard error mean (SEM). Bar graphs represent mean + SEM. ****p < 0.0001, **p < 0.01 ordinary two-way ANOVA comparing to cells treated with DMSO. Scale bars = 10µm.

**Supplementary Table 1:** LOCKR-S candidates tested. RBD-LOCKR-S in Tab 1 or Ras-LOCKR-S in Tab 2 that were tested in this study. Domain structure, sequence, and performance to either RBD expression (RBD-LOCKR-S) or EGF stimulation (Ras-LOCKR-S) are documented.

**Supplementary Table 2:** LOCKR-PL candidates tested. RBD-LOCKR-PL in Tab 1 or Ras-LOCKR-PL in Tab 2 that were tested in this study. Domain structure, sequence, and performance to either RBD expression (RBD-LOCKR-PL) or EGF stimulation (Ras-LOCKR-PL) are documented.

**Supplementary Table 3:** Mass spectrometry data of Ras-active EML4-Alk granules. Some Beas2B cells transfected with EML4-Alk v1 or v3 plus GFPnb-Ras-LOCKR-PL. Some Beas2B cells were transfected with GFPnb-Ras-LOCKR-PL alone. EML4-Alk v1/v3 plus GFPnb-Ras-LOCKR-PL compared to GFPnb-Ras-LOCKR-PL. Different biotin incubation lengths were considered.

**Supplementary Table 4:** Resources and identifiers. Details of cells, molecules, and software used in this study.

## Notes

### Summary of Updates

Manuscript has been streamlined to focus on the development of Ras sensors.

## References

1. Cox, A. D. & Der, C. J. Ras history. Small GTPases 1, 2–27 (2010).

2. Augsten, M. et al. Live-cell imaging of endogenous Ras-GTP illustrates predominant Ras activation at the plasma membrane. EMBO Rep 7, 46–51 (2006).

3. Bivona, T. G. et al. PKC regulates a farnesyl-electrostatic switch on K-Ras that promotes its association with Bcl-XL on mitochondria and induces apoptosis. Mol Cell 21, 481–493 (2006).

4. Chiu, V. K. et al. Ras signalling on the endoplasmic reticulum and the Golgi. Nat Cell Biol 4, 343–350 (2002).

5. Tulpule, A. et al. Kinase-mediated RAS signaling via membraneless cytoplasmic protein granules. Cell 184, 2649–2664.e18 (2021).

6. Rose, J. C. et al. A computationally engineered RAS rheostat reveals RAS-ERK signaling dynamics. Nat Chem Biol 13, 119–126 (2017).

7. Bivona, T. G. et al. Phospholipase Cγ activates Ras on the Golgi apparatus by means of RasGRP1. Nature 424, 694–698 (2003).

8. Komatsu, N. et al. Development of an optimized backbone of FRET biosensors for kinases and GTPases. Mol Biol Cell 22, 4647–4656 (2011).

9. Mochizuki, N. et al. Spatio-temporal images of growth-factor-induced activation of Ras and Rap1. Nature 411, 1065–1068 (2001).

10. Yang, H. H. & St-Pierre, F. Genetically encoded voltage indicators: opportunities and challenges. Journal of Neuroscience 36, 9977–9989 (2016).

11. Li, Y.-C., et al. Analysis of RAS protein interactions in living cells reveals a mechanism for pan-RAS depletion by membrane-targeted RAS binders. Proceedings of the National Academy of Sciences 117, 12121–12130 (2020).

12. Sanford, L. & Palmer, A. Recent advances in development of genetically encoded fluorescent sensors. Methods Enzymol 589, 1–49 (2017).

13. Langan, R. A. et al. De novo design of bioactive protein switches. Nature 572, 205–210 (2019).

14. Quijano-Rubio, A., et al. De novo design of modular and tunable protein biosensors. Nature (2021) doi:10.1038/s41586-021-03258-z.

15. Greenwald, E. C., Mehta, S. & Zhang, J. Genetically Encoded Fluorescent Biosensors Illuminate the Spatiotemporal Regulation of Signaling Networks. Chem Rev 118, 11707– 11794 (2018).

16. Zhang, J. Z. et al. Thermodynamically coupled biosensors for detecting neutralizing antibodies against SARS-CoV-2 variants. Nat Biotechnol (2022) doi:10.1038/s41587-022-01280-8.

17. Tran, T. H. et al. KRAS interaction with RAF1 RAS-binding domain and cysteine-rich domain provides insights into RAS-mediated RAF activation. Nat Commun 12, 1176 (2021).

18. Jumper, J. et al. Highly accurate protein structure prediction with AlphaFold. Nature 596, 583–589 (2021).

19. O’Shaughnessy, E. C. et al. Software for lattice light-sheet imaging of FRET biosensors, illustrated with a new Rap1 biosensor. Journal of Cell Biology 218, 3153–3160 (2019).

20. Zhou, X. et al. Location-specific inhibition of Akt reveals regulation of mTORC1 activity in the nucleus. Nat Commun 11, 6088 (2020).

21. Tenner, B., Zhang, J. Z., Huang, B., Mehta, S. & Zhang, J. FluoSTEPs: Fluorescent biosensors for monitoring compartmentalized signaling within endogenous microdomains. Sci Adv 7, eabe4091 (2021).

22. Gao, X. & Zhang, J. Spatiotemporal analysis of differential Akt regulation in plasma membrane microdomains. Mol Biol Cell 19, 4366–4373 (2008).

23. Fulton, D. et al. Targeting of endothelial nitric-oxide synthase to the cytoplasmic face of the Golgi complex or plasma membrane regulates Akt- versus calcium-dependent mechanisms for nitric oxide release. J Biol Chem 279, 30349–30357 (2004).

24. Choy, E. et al. Endomembrane trafficking of ras: the CAAX motif targets proteins to the ER and Golgi. Cell 98, 69–80 (1999).

25. Quatela, S. E. & Philips, M. R. Ras signaling on the Golgi. Curr Opin Cell Biol 18, 162–167 (2006).

26. Mor, A. et al. The lymphocyte function-associated antigen-1 receptor costimulates plasma membrane Ras via phospholipase D2. Nat Cell Biol 9, 713–719 (2007).

27. Kwak, C. et al. Contact-ID, a tool for profiling organelle contact sites, reveals regulatory proteins of mitochondrial-associated membrane formation. Proceedings of the National Academy of Sciences 117, 12109–12120 (2020).

28. Cho, K. F. et al. Split-TurboID enables contact-dependent proximity labeling in cells. Proceedings of the National Academy of Sciences 117, 12143 LP – 12154 (2020).

29. Keyes, J. et al. Signaling diversity enabled by Rap1-regulated plasma membrane ERK with distinct temporal dynamics. Elife 9, (2020).

30. Sasaki, T., Rodig, S. J., Chirieac, L. R. & Jänne, P. A. The biology and treatment of EML4- ALK non-small cell lung cancer. Eur J Cancer 46, 1773–1780 (2010).

31. Sampson, J., Richards, M. W., Choi, J., Fry, A. M. & Bayliss, R. Phase-separated foci of EML4-ALK facilitate signalling and depend upon an active kinase conformation. EMBO Rep 22, e53693 (2021).

32. Arkun, Y. & Yasemi, M. Dynamics and control of the ERK signaling pathway: Sensitivity, bistability, and oscillations. PLoS One 13, e0195513 (2018).

33. Sabbatini, M. E. & Williams, J. A. Cholecystokinin-mediated RhoGDI phosphorylation via PKCα promotes both RhoA and Rac1 signaling. PLoS One 8, e66029–e66029 (2013).

34. Tnimov, Z. et al. Quantitative analysis of prenylated RhoA interaction with its chaperone, RhoGDI. J Biol Chem 287, 26549–26562 (2012).

35. Barlat, I. et al. A role for Sam68 in cell cycle progression antagonized by a spliced variant within the KH domain. Journal of Biological Chemistry 272, 3129–3132 (1997).

36. Bielli, P., Busà, R., Paronetto, M. P. & Sette, C. The RNA-binding protein Sam68 is a multifunctional player in human cancer. Endocr Relat Cancer 18, R91–R102 (2011).

37. Masibag, A. N. et al. Pharmacological targeting of Sam68 functions in colorectal cancer stem cells. iScience 24, 103442 (2021).

38. Rawlings, D. J., Sommer, K. & Moreno-García, M. E. The CARMA1 signalosome links the signalling machinery of adaptive and innate immunity in lymphocytes. Nat Rev Immunol 6, 799–812 (2006).

39. Werlen, G. & Palmer, E. The T-cell receptor signalosome: a dynamic structure with expanding complexity. Curr Opin Immunol 14, 299–305 (2002).

40. Karniol, B. & Chamovitz, D. A. The COP9 signalosome: from light signaling to general developmental regulation and back. Curr Opin Plant Biol 3, 387–393 (2000).

41. Yablonski, D. Bridging the gap: modulatory roles of the Grb2-family adaptor, Gads, in cellular and allergic immune responses. Front Immunol 10, 1704 (2019).

42. Huang, W. Y. C. et al. A molecular assembly phase transition and kinetic proofreading modulate Ras activation by SOS. Science (1979) 363, 1098 LP – 1103 (2019).

43. Xiong, Z. et al. In vivo proteomic mapping through GFP-directed proximity-dependent biotin labelling in zebrafish. Elife 10, e64631 (2021).

44. Zhang, J. Z. et al. Histamine-induced biphasic activation of RhoA allows for persistent RhoA signaling. PLoS Biol 18, e3000866 (2020).

45. Scrima, A., Thomas, C., Deaconescu, D. & Wittinghofer, A. The Rap-RapGAP complex: GTP hydrolysis without catalytic glutamine and arginine residues. EMBO J 27, 1145–1153 (2008).

46. Satoh, A. et al. The Golgin Protein Giantin Regulates Interconnections Between Golgi Stacks. Front Cell Dev Biol 7, (2019).

